# Genetic analysis argues for a coactivator function for the *Saccharomyces cerevisiae* Tup1 corepressor

**DOI:** 10.1101/2021.05.11.443680

**Authors:** Emily J. Parnell, Timothy J. Parnell, David J. Stillman

## Abstract

The Tup1-Cyc8 corepressor complex of *Saccharomyces cerevisiae* is recruited to promoters by DNA-binding proteins to repress transcription of genes, including the **a**-specific mating type genes. We report here a *tup1(S649F)* mutant that displays mating irregularities similar to a *tup1* null and an α-predominant growth defect. RNA-Seq and ChIP-Seq were used to analyze gene expression and Tup1 occupancy changes in mutant vs. wild-type in both **a** and α cells. Increased Tup1(S649F) occupancy tended to occur upstream of upregulated genes, whereas locations with decreased occupancy usually did not show changes in gene expression, suggesting this mutant not only loses corepressor function but also behaves as a coactivator. Based upon studies demonstrating a dual role of Tup1 in both repression and activation, we postulate that the coactivator function of Tup1(S649F) results from diminished interaction with repressor proteins, including α2. We also found that large changes in mating type-specific gene expression between **a** and α or between mutant and wild-type were not easily explained by the range of Tup1 occupancy levels within their promoters, as predicted by the classic model of **a**-specific gene repression by Tup1. Most surprisingly, we observed Tup1 occupancy upstream of the **a**-specific gene *MFA2* and the α-specific gene *MF(ALPHA)1* in cells in which each gene was expressed rather than repressed. These results, combined with identification of additional mating related genes upregulated in the *tup1(S649F)* α strain, illustrate that the role of Tup1 in distinguishing mating types in yeast appears to be both more comprehensive and more nuanced than previously appreciated.

## INTRODUCTION

Regulation of gene expression at complex promoters is facilitated by a host of factors, including sequence specific DNA-binding factors and proteins that are recruited as coactivators or corepressors. Some of these complexes, including SAGA and Sin3/Rpd3, modify the amino-terminal tails of histones to promote either activation or repression (Wolffe 1996; Ng and Bird 2000; Verdone *et al*. 2005; Shahbazian and Grunstein 2007; Soffers and Workman 2020). Others are nucleosome remodelers that move or evict nucleosomes, such as SWI/SNF and Isw2 (Lin *et al*. 2020; Markert and Luger 2021; Reyes *et al*. 2021). A balance of activating and repressing functions is required to maintain appropriate gene expression levels, and increasing evidence points to an integration of these two processes. Some multi-protein complexes can affect transcription either positively or negatively, suggesting their roles may be context dependent rather than fitting within strict categories (Kliewe *et al*. 2017; Adams *et al*. 2018).

The Tup1-Cyc8 complex of *Saccharomyces cerevisiae* was one of the first to be identified as a transcriptional corepressor (Keleher *et al*. 1992; Tzamarias and Struhl 1994). This 1.2 megadalton complex consists of a 4:1 ratio of Tup1 to Cyc8 (Williams *et al*. 1991; Varanasi *et al*. 1996). Neither protein binds DNA directly, but when tethered upstream of a heterologous reporter, Tup1 and Cyc8 can confer repression (Tzamarias and Struhl 1994). Tup1-Cyc8 is recruited to native promoters by DNA-binding proteins, thereby repressing diverse groups of genes including cell-type specific genes (**a** haploid vs. α haploid vs. diploid), glucose- and hypoxia-regulated genes, as well as genes that respond to osmotic stress or DNA damage and genes involved in flocculation (Lipke and Hull-Pillsbury 1984; Malave and Dent 2006). In total, Tup1-Cyc8 affects the expression of approximately 300-500 genes in budding yeast (Derisi *et al*. 1997; Green and Johnson 2004; Chen *et al*. 2013). Both Tup1 and Cyc8 have sequence homologues in *Schizosacchaomyces pombe* and *Caenorhabditis elegans* as well as evolutionary relatives in higher eukaryotes, such as Groucho in *Drosophila melanogaster* and TLE1 in humans (Chen and Courey 2000). Metazoan members of the Groucho/TLE family of corepressors have similar domain structures and corepressor functions as Tup1 and regulate genes that are essential to many developmental pathways or are implicated in various cancers (Fisher and Caudy 1998; Grbavec *et al*. 1998; Parkhurst 1998; Grbavec *et al*. 1999; Chen and Courey 2000).

The C-terminus of Tup1 folds into a WD domain with seven repeats that form the blades of a propeller-like structure (Williams and Trumbly 1990). This protein-protein interaction domain binds to the homeodomain-containing protein α2, the repressor of **a**-specific genes via α2/Mcm1 and of haploid-specific genes via **a**1/α2 (Johnson and Herskowitz 1985; Keleher *et al*. 1989; Dranginis 1990; Komachi *et al*. 1994). Deletion of the WD domain is effectively a *tup1* null, and mutations within the WD domain affect target genes other than the α2-regulated genes, suggesting it interacts with additional DNA-binding factors or proteins required for repression (Williams and Trumbly 1990; Carrico and Zitomer 1998). The N-terminus of Tup1 facilitates its tetramerization as well as interaction of tetramers with Cyc8 (Jabet *et al*. 2000). The central portion of the protein contains two additional domains with repressive functions, with the more C-terminal domain playing a more substantial role (Tzamarias and Struhl 1994; Zhang *et al*. 2002). Cyc8 has 10 tandem repeats of a tetratricopeptide repeat (TPR) sequence motif, capable of interacting with diverse proteins, including Tup1 (Smith *et al*. 1995; Tzamarias and Struhl 1995; Gounalaki *et al*. 2000). The combination of Tup1 and Cyc8 thus provides ample diverse surfaces for interaction with proteins, allowing them to be recruited in a range of different contexts to affect the expression of a large number of genes.

A number of mechanisms for Tup1-Cyc8 repression have been proposed, including recruitment of histone deacetylases, formation of compact nucleosomal arrays, as well as interference with the Mediator complex and/or RNA Polymerase II itself (Smith and Johnson 2000; Malave and Dent 2006). These possibilities are not mutually exclusive, and it seems likely that distinct, possibly redundant mechanisms operate at different promoters (Zhang and Reese 2004). A potential unifying model for Tup1-Cyc8 repression suggests that it masks the activation domains of its many DNA-binding recruiter proteins, thereby preventing association of many recruiters with coactivator proteins (Wong and Struhl 2011). In this model, the DNA-binding proteins that associate with Tup1-Cyc8 would be capable of recruiting both coactivators and the Tup1-Cyc8 corepressor. Environmental conditions could prompt release from the repressed, masked state, possibly by a postranslational modification or a conformational change of either the DNA-binding protein or Tup1-Cyc8.

Several studies support the conclusion that Tup1-Cyc8 can function as more than a simple corepressor. Tup1 associates with some of its target promoters during gene activation as well as during repression, in some cases even facilitating recruitment of coactivators such as SAGA and SWI/SNF (Papamichos-Chronakis *et al*. 2002; Proft and Struhl 2002; Mennella *et al*. 2003; Desimone and Laney 2010). Tup1’s continued presence throughout induction may be important for priming these promoters for subsequent reactivation following repression. At some glucose-repressed genes, Tup1 is responsible for Htz1 deposition at the promoter proximal nucleosome, a mark that is needed for rapid recruitment of Mediator and activation following Tup1-Cyc8-mediated repression (Gligoris *et al*. 2007). Similarly, at the repressed promoters of mating-type-specific genes, Gcn5 acetylation of histone H3 amino-terminal tails requires Tup1-Cyc8 (Desimone and Laney 2010). In these cases, Tup1 appears to repress transcription while also providing a chromatin state that allows for a rapid switch back to activation. A coactivator function of Tup1 has been observed at additional genes, including those encoding tryptophan and other amino acid transporters, the *BAP2* amino acid permease gene, the *CIT2* citrate synthase gene, the *FRE2* plasma membrane ferric reductase gene, and the *FLO11* flocculation gene (Conlan *et al*. 1999; Nielsen *et al*. 2001; Fragiadakis *et al*. 2004; Tanaka and Mukai 2015; Nguyen *et al*. 2018).

The picture emerging from studies of Tup1-Cyc8 is that the complex would be most aptly termed a transcriptional “coregulator”. The persistence of Tup1-Cyc8 association as promoters are activated following repression suggests there is a functional switch that converts the complex from a corepressor to a coactivator. Understanding these two functions of Tup1-Cyc8 is complicated by the fact that the complex appears to play different roles at subsets of promoters. It has also become increasingly clear that Tup1-Cyc8 target promoters have multiple, redundant mechanisms of recruitment, with binding sites for distinct DNA-binding proteins that are each capable of recruiting Tup1 and/or Cyc8. Using different combinations of these binding sites, promoters are able to repress transcription in response to different environmental conditions (Hanlon *et al*. 2011; Tam and Van Werven 2020). Some Tup1-Cyc8 association sites cannot be explained by the current list of recruiter proteins, suggesting that additional DNA-binding proteins that recruit Tup1-Cyc8 have yet to be identified.

We recently identified the Ash1 DNA-binding repressor protein as an additional Tup1 recruiter (Parnell *et al*. 2020). Ash1 is necessary for daughter cell-specific repression of the *HO* gene, which encodes the endonuclease that cleaves the *MAT* locus to allow haploid yeast cells to switch mating type (Strathern *et al*. 1982; Bobola *et al*. 1996; Sil and Herskowitz 1996). The *HO* promoter is highly regulated, utilizing multiple coactivators and corepressors. Recruitment of Tup1 to *HO* by Ash1 occurs in an activator-dependent manner and may be important for resetting the promoter for repression following a brief burst of gene activation at the end of the G1 phase of the cell cycle (Parnell *et al*. 2020). Ash1 colocalizes with Tup1 at a number of other genomic sites, suggesting it recruits Tup1 to additional promoters.

Our discovery of Tup1 repression of the *HO* promoter in haploid cells resulted from a genetic screen for negative regulators of the *HO* promoter (Parnell and Stillman 2019). In this screen, we identified a *tup1* hypomorph, *tup1(H575Y)*, which allowed some expression of *HO* in the absence of its pioneer transcription factor, Swi5. Here we report new, targeted screens to identify additional *tup1* alleles, with the goal of better understanding the role of Tup1, both at the *HO* promoter and genome-wide. RNA-Seq and ChIP-Seq experiments with the *tup1(S649F)* mutant identified in this new screen provide further evidence that Tup1 has a coactivator as well as a corepressor function and that this dichotomy of function is not restricted to a small number of genes but is a general feature of Tup1 regulation. Our study also provides new insight into the role of Tup1 at the mating type-specific genes, some of the prototypical genes that first suggested the corepressor function of the Tup1-Cyc8 complex.

## RESULTS

### Genetic screens identified several Tup1 WD domain mutants that increase *HO* expression

Having identified a *tup1* mutation as a suppressor of *HO* transcription in the absence of the Swi5 transcription factor (Parnell and Stillman 2019), we next sought to identify additional residues of Tup1 that participate in repression of the *HO* promoter by conducting two targeted genetic screens. Each screen was performed using a yeast strain in which *HO* expression was decreased due to the absence of an activator or coactivator of the *HO* promoter. Strains carried an *HO-ADE2* reporter, allowing the level of *HO* expression to be observed as growth on media lacking adenine (Jansen *et al*. 1996). The chromosomal copy of *TUP1* was deleted and covered with a *TUP1*-containing plasmid. We then swapped this with a PCR-mutagenized version of *TUP1* and isolated alleles that elevated *HO-ADE2* expression, as determined by increased growth on -Ade media.

The first screen with the mutagenized *TUP1* gene was conducted in a *swi5 ash1* strain similar to that used to identify the *tup1(H575Y)* allele (Parnell and Stillman 2019). Swi5 is the pioneer activator of *HO* expression; its binding to two sites within the promoter sets in motion a series of factor recruitments, histone-modifications, and nucleosome evictions that ultimately allow activation of transcription (Cosma *et al*. 1999; Bhoite *et al*. 2001; Takahata *et al*. 2009; Stillman 2013). Expression of *HO-ADE2* in a *swi5* mutant is extremely low, and cells grow very poorly in absence of adenine (Parnell and Stillman 2019). Ash1 is a repressor of *HO* expression that acts predominantly in daughter cells, leading to expression of *HO* only in mother cells (Bobola *et al*. 1996; Sil and Herskowitz 1996). We previously determined that additional negative regulators of *HO* could be detected using the *HO-ADE2* reporter, but only when an *ash1* mutation was added to the *swi5* strain. This increased the baseline expression of the *HO* promoter enough to allow loss of another repressor to be visible as increased growth on media lacking adenine (Parnell and Stillman 2019). Several negative transcription factors participate in *HO* regulation, and substantial increases in expression were only observed upon removal of multiple of these factors simultaneously (Parnell and Stillman 2019).

While the screen conducted in a *swi5 ash1* strain was productive for isolating the *tup1(H575Y)* mutant allele, the later discovery that Ash1 is responsible for most of the Tup1 recruitment to the *HO* promoter suggests an *ASH1* strain could be better suited for identification of *tup1* alleles (Parnell *et al*. 2020). Therefore, we conducted a second screen involving a mutagenized *TUP1* plasmid in a *gcn5* mutant background. Gcn5 is the catalytic subunit of a histone acetyltransferase complex required for full *HO* expression (Cosma *et al*. 1999; Takahata *et al*. 2011). In the absence of Gcn5, *HO-ADE2* expression decreases to a level similar to that of a *swi5 ash1* strain, providing a baseline level of growth on -Ade media that allowed small increases in *HO-ADE2* expression to be visible as more robust growth (Parnell and Stillman 2019).

Mutants of Tup1 identified in the two screens are listed in Table 1. The apparent bias toward mutations identified in the *swi5 ash1* screen reflects different validation procedures used for the screens rather than a difference in the sensitivity of the screens (see Methods for additional explanation). While the entire *TUP1* gene was PCR-mutagenized, all mutants from the two screens fell within the WD domain of the protein, known to interact with the α2 repressor and suggested to participate in multiple types of protein-protein interactions with other DNA-binding proteins (Komachi *et al*. 1994; Carrico and Zitomer 1998).

**Table 1.** Tup1 mutants obtained from genetic screens for activation of *HO* expression.

| AA Position | WT AA | Mutant AA | # Indep Mutants | Classification<sup>a</sup> | Screen | Position within WD | Structural Information |
| --- | --- | --- | --- | --- | --- | --- | --- |
| 445 | Y | C | 2 | 3 | *gcn5* | DA Loop 1-2 | Top Surface Previously identified |
| 526<sup>b</sup> | E | G | 2 | 3 | *swi5 ash1* | DA Loop 3-4 | Top Surface |
| 560 | F | S, Y | 2 | 3 | *swi5 ash1* | CD Loop 4<sup>th</sup> | Side Surface |
| 564 | R | G | 2 | 3 | *swi5 ash1* | D Sheet 4<sup>th</sup> | Side Surface |
| 565 | L | P | 2 | 1,2 | *swi5 ash1* | D Sheet 4<sup>th</sup> | Just Under Side Surface |
| 566 | D | G | 3 | 1,2 | *swi5 ash1* | DA Loop 4-5 | Side Surface |
| 569 | N | D, K | 2 | 3 | *swi5 ash1* | DA Loop 4-5 | Not in structure |
| 570 | E | G, K, V | 6 | 3 | *swi5 ash1* | DA Loop 4-5 | Not in structure |
| 575 | H | P, R, Y | 7 | 1,2 | *swi5 ash1, gcn5* | DA Loop 4-5 | Just Under Top Surface |
| 597 | D | G, N, Y | 12 | 1,2 | *swi5 ash1* | BC Turn 5<sup>th</sup> | Top Surface |
| 602 | L | P | 2 | 3 | *gcn5* | C Sheet 5<sup>th</sup> |  |
| 629 | H | R, Y | 2 | 3 | *swi5 ash1, gcn5* | DA Loop 5-6 | Just Under Top Surface |
| 649 | S | F | 2 | 3 | *gcn5* | B Sheet 6<sup>th</sup> | Just Under Top Surface |
| 673<sup>b</sup> | N | D | 15 | 3 | *swi5 ash1* | DA Loop 6-7 | Top Surface Previously identified |
| 699 | D | G | 3 | 1,2 | *swi5 ash1, gcn5* | BC Turn 7<sup>th</sup> | Top Surface |
| 700 | C | R, Y | 2 | 1,2 | *swi5 ash1* | BC Turn 7<sup>th</sup> | Top Surface |

<sup>a</sup> Classification designation are as follows:
1 = Single mutation; verified by re-transformation of mutant plasmid and confirmation of phenotype.
2 = Mutation position is identical to a single, verified mutation. However, this mutant either has additional positions that are mutated or was not verified.
3 = Mutation was identified in more than one mutant, but none are both single and verified.
<sup>b</sup> Mutation position is identical to one previously identified by Komachi et al (MCB 1997).
Red font indicates alleles that were constructed at the endogenous locus and tested in Fig 2.

Most mutations were located within WD repeats 4-7, including multiple hits at position H575, which was the site of the originally isolated *tup1(H575Y)* allele (Table 1, Fig 1A, B). Several mutations were also identified within the DA loops between successive WD repeats (Table 1; Y445, E526, N569, E570, H575, H629, N673). Of these residues, Y445, E526 and N673 are exposed on the top surface of the WD domain (Fig 1B). Y445 and N673 were previously identified as contact points with the α2 repressor (Komachi *et al*. 1994).

**Figure 1.**
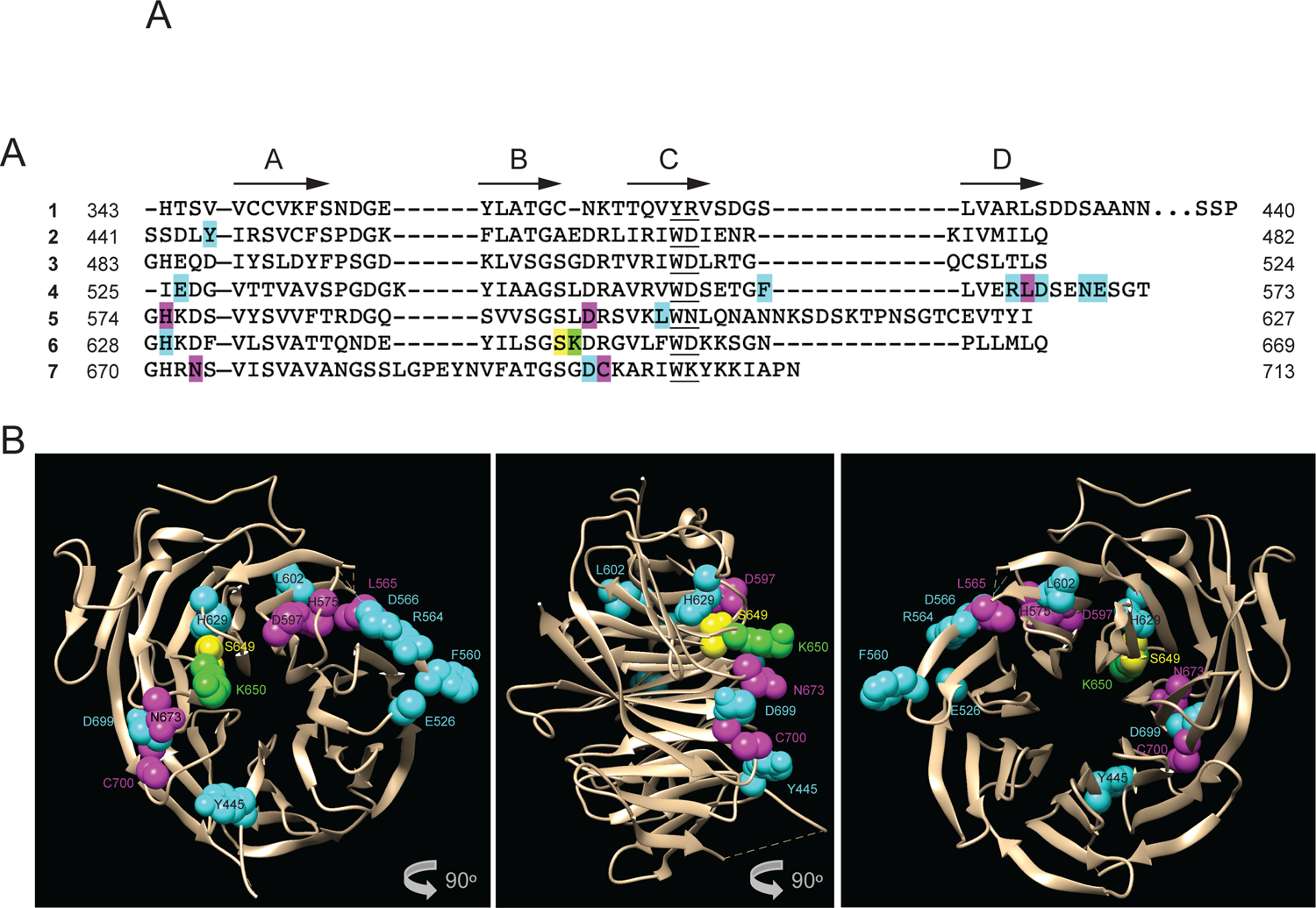
Tup1 mutants identified in genetic screens for activation of *HO* expression. (A) All *tup1* mutations were located with the WD domain at the C-terminus of Tup1. The amino acid sequence of the WD domain of Tup1 is shown, with each line representing the WD repeat number on the left. Amino acid number positions are indicated. Beta sheets determined in Sprague et al (2000) are indicated by arrows and the A-D designation above the sequence. Forty-eight amino acids from the DA loop between WD repeats 1 and 2 were eliminated for simplicity (H390 to S437), as indicated by “…”. The WD or variant motif within each repeat is underlined. Highlighted amino acids represent mutation positions: yellow for S649, green for K650, magenta for the other five mutants for which endogenous mutants were constructed and tested for effects on *HO* expression (L565, D597, N673, C700), and teal for all other mutants listed in Table 1. (B) Most mutations lie on one surface of the propeller. Structure of the WD domain of Tup1 (Sprague *et al*. 2000) is shown, with mutant positions indicated. Color coding is the same as in A. Note that mutants at positions N569 and E570 are not shown because they were not in the resolved structure. The central image is a 90° rotation relative to the left image, to show the amino acids, including K650, that project from the surface of Tup1, and the right image is another 90° rotation to illustrate the opposite surface.

Multiple sites affected by the mutations are likely important for the stability of their respective WD repeats. Within each repeat, interactions among a structural tetrad of amino acids are important for maintaining stability; these include the tryptophan in the C sheet, the serine/threonine in the B sheet, the histidine in the DA loop and the aspartic acid in the tight BC turn (Sprague *et al*. 2000). Our screens identified mutations in S649, H575, H629, D597 and D699, all of which are residues thought to participate in these structural interactions. The aspartic acid within this structural tetrad is the most invariant amino acid position across WD repeats in different proteins, and two of our mutations are at these locations (D597 and D699) (Sprague *et al*. 2000). The importance of these positions is also highlighted by the observation that mutations in positions H575, H629 and D699 were identified in both screens.

### The *tup1(S649F)* mutant displays an α cell-predominant slow growth phenotype

We chose a panel of six mutations for further study, representing the most frequently mutated amino acids and a variety of positions within the WD domain (Table 1, indicated in red). These include the original *tup1(H575Y)* allele as well as *tup1(L565P)*, *tup1(D597N)*, *tup1(S649F)*, *tup1(N673D)*, and *tup1(C700R)*. Strains were constructed with each of these mutations at the endogenous, unmarked *TUP1* locus and then assessed for phenotypes resembling those of *tup1* null mutants.

Strains that are *tup1* null grow poorly and experience a delay in G1 of the cell cycle. An accurate assessment of viability of *tup1* null strains in a growth assay is difficult to obtain due to the frequent occurrence of suppressors, at least in the W303 background. Of the *tup1* point mutants we examined, only *tup1(S649F)* showed an obvious slow growth phenotype at the three temperatures examined (Fig 2). While the effect was subtle in the *tup1(S649F) MAT***a** strain, there was a more obvious growth defect at all temperatures in the *MAT*α strain.

**Figure 2.**
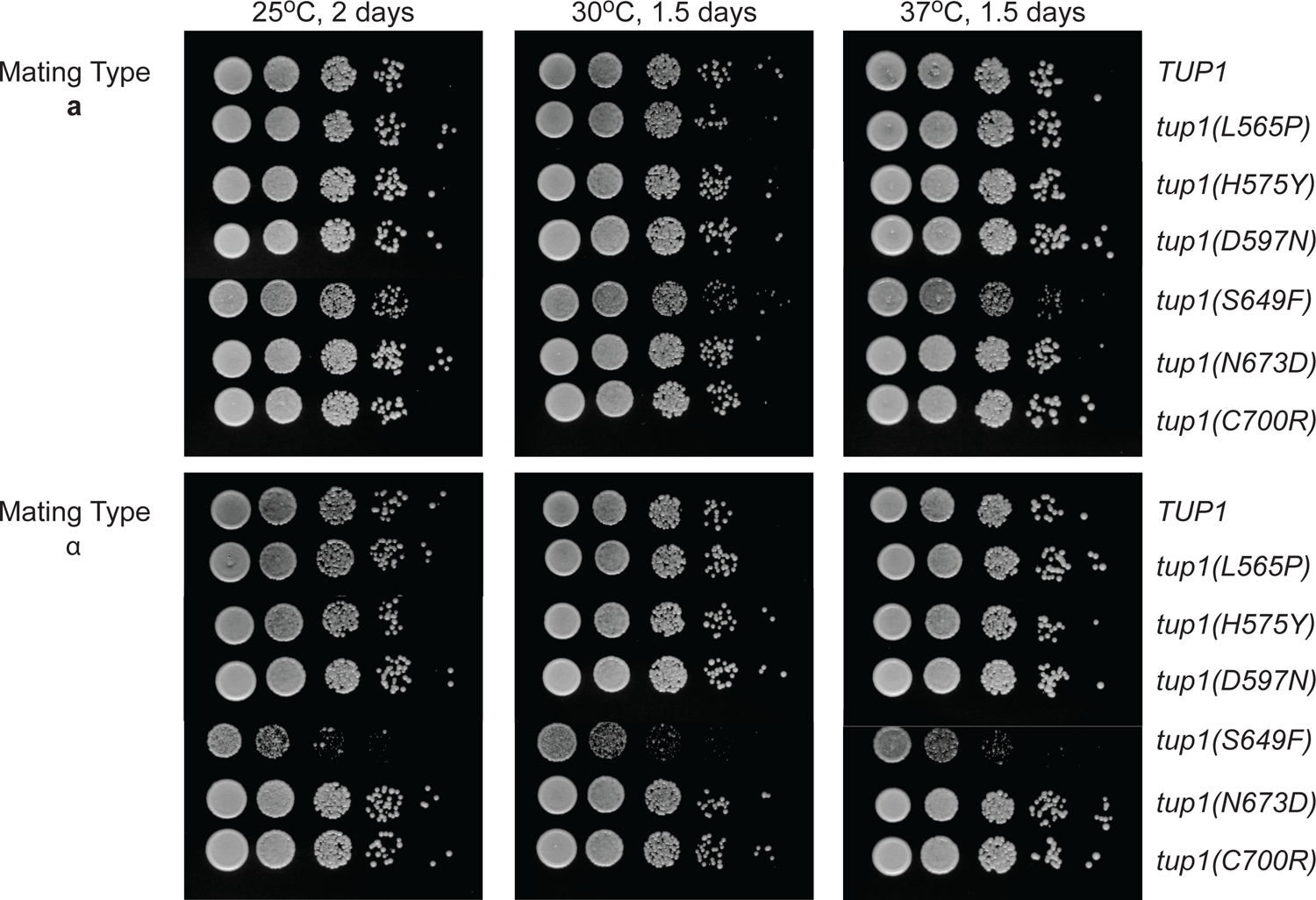
The *tup1(S649F)* mutant shows a growth defect that is stronger in the α strain. Shown are 10-fold serial dilutions of the *tup1* mutants indicated at the right, compared to *TUP1* wild-type, grown on YPAD media at the indicated temperatures and time durations. Strains with mating type **a** are on top; strains with mating type α are on the bottom.

MATα *tup1* strains show decreased mating to **a** strains, increased mating to α strains, and a “shmoo”-like phenotype reminiscent of **a** cells that are responding to α-factor, resulting from mis-regulation of mating type-specific genes normally repressed by Tup1 (Wickner 1974; Lemontt *et al*. 1980; Mukai *et al*. 1991). The *tup1(S649F)* α cells also had a shmoo-like appearance and experienced a delay in mating but later was capable of mating with both **a** and α *TUP1* strains (data not shown). Other *tup1* point mutants did not display mating irregularities (data not shown).

Cells lacking *TUP1* are also flocculent, due to inappropriate expression of the agglutinin genes, a class of cell surface glycoproteins repressed by Tup1 (Lipke and Hull-Pillsbury 1984). Most *tup1* mutant alleles had only minor flocculation phenotypes (data not shown). However, both **a** and α *tup1(S649F)* strains displayed a strong flocculation phenotype, with the effect in the α strain more severe than in the **a** strain, approaching the level of flocculation of a *tup1* null. Collectively, the reduced growth, mating characteristics and flocculation phenotype of *tup1(S649F)* suggest this mutant substantially diminishes but does not eliminate Tup1 function.

### The *tup1(S649F)* mutant allows substantial increases in *HO* expression

To assess the effects of the *tup1* point mutants on native *HO* expression rather the *HO-ADE2* reporter used for our screens, we measured the level of endogenous *HO* expression in *tup1* mutants by RT-qPCR within the context of a *swi5 ash1* strain, a *gcn5* strain and an otherwise wild-type strain (Fig 3A, B, C). All *tup1* mutations increased expression of *HO* in the *swi5 ash1* strain, even the *tup1(S649F)* mutation isolated only from the *gcn5* screen (Fig 3A). The *tup1(S649F)* and *tup1(N673D)* mutations increased expression most strongly, more than 3-fold over the level of *HO* expression in the *swi5 ash1* strain. All mutations also increased *HO* expression in a *gcn5* strain with a significance level of *p* < 0.05, but most of these increases were less than two-fold (Fig 3B). The exceptions were *tup1(D597N)* (just over two-fold increase) and *tup1(S649F),* identified originally only in the *gcn5* screen (nearly 7-fold increased expression). Within the context of an otherwise wild-type strain, most mutations increased *HO* expression only very modestly (Fig 3C). The *tup1(S649F)* mutation, however, displayed a greater than 2-fold increase in expression.

**Figure 3.**
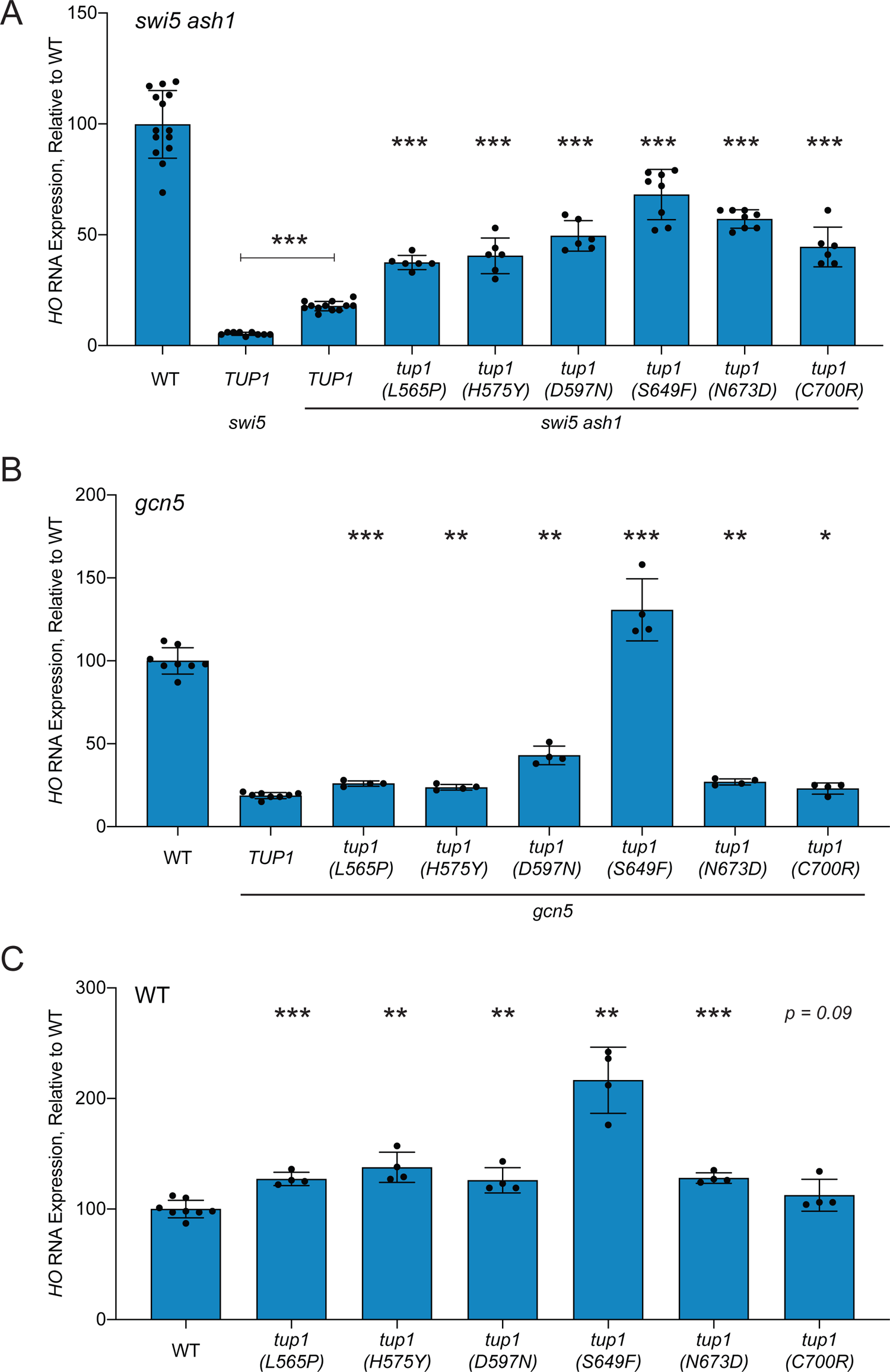
*HO* expression increases substantially in the *tup1(S649F)* mutant. *HO* mRNA levels were measured by RT-qPCR, normalized to *RPR1*, and expressed relative to wild-type. Each dot represents a single data point, and error bars reflect the standard deviation. *** *p* < 0.001, ** *p* < 0.01, * *p* < 0.05. (A) Mutant *tup1* alleles increase *HO* expression in a *swi5 ash1* strain. (B) The *tup1(S649F)* mutant displays strong suppression of a *gcn5* mutant at the *HO* gene. (C) The *tup1(S649F)* mutant increases *HO* expression in an otherwise wild-type context.

Our genetic screens were therefore effective in identifying additional *tup1* mutations at a variety of positions in the WD domain that suppress *swi5 ash1* and *gcn5* for expression of *HO* and likely affect regulation of additional Tup1 target genes. The strongest of these was *tup1(S649F)*, as measured by its effect on growth, mating, flocculation and *HO* expression both alone and via suppression of *swi5 ash1* and *gcn5* mutants. The S649 amino acid has been predicted to contribute to structural stability of the 6th WD repeat via a tetrad of interactions between H629, S649, D651 and W657 (Sprague 2000). Loss of integrity of this repeat could destabilize the WD domain and possibly the entire protein. To examine this possibility, we tagged wild-type Tup1 and Tup1(S649F) with a V5 epitope tag and visualized protein expression and stability by western blot (Fig S1). As predicted, the protein level of Tup1(S649F) was reduced relative to wild-type. Effects of the *tup1(S649F)* mutation could therefore largely result from a diminished level of stable protein.

### Effects of *tup1(S649F)* on *HO* transcription are not caused by a loss of post-translational modifications

An alternative possibility for the effects of the *tup1(S649F)* mutation on growth and *HO* expression could be a loss of post-translational modification of Tup1 via one of two mechanisms. Serine 649 could be modified by phosphorylation, or the immediately adjacent amino acid, a lysine residue at position 650, could be acetylated. Residue K650 appears to project outward from one surface of the WD domain, and S649 is positioned just beneath it (Fig 1B), suggesting that the effect of *tup1(S649F)* may be the introduction of a bulky amino acid that would alter the positioning of K650, impairing the ability to be acetylated or deacetylated. There is precedent in the literature for regulation of chromatin and transcriptional-related proteins by acetylation (Narita *et al*. 2019). One of these methods of post-translational modification could provide a mechanism for switching between active and repressive forms of Tup1, consistent with the model in which Tup1 masks activation domains but subsequently participates in gene activation. We tested these possibilities by constructing additional mutants at positions 649 and 650 and measuring suppression of *HO* expression in *swi5 ash1* and *gcn5* strains relative to the original *tup1(S649F)* mutant.

A *tup1(S649A)* mutant was used to test the possibility that an inability to phosphorylate S649 is the critical reason for the effect of *tup1(S649F)* on *HO* expression. The *tup1(S649A)* mutation, in sharp contrast to *tup1(S649F)*, had no effect on *HO* expression in either a *swi5 ash1* or a *gcn5* strain (Fig 4A, 4B). This result demonstrates that introduction of the phenylalanine amino acid, rather than a simple loss of possible S649 phosphorylation by substitution of a different amino acid, is important for the observed *tup1(S649F)* phenotype.

**Figure 4.**
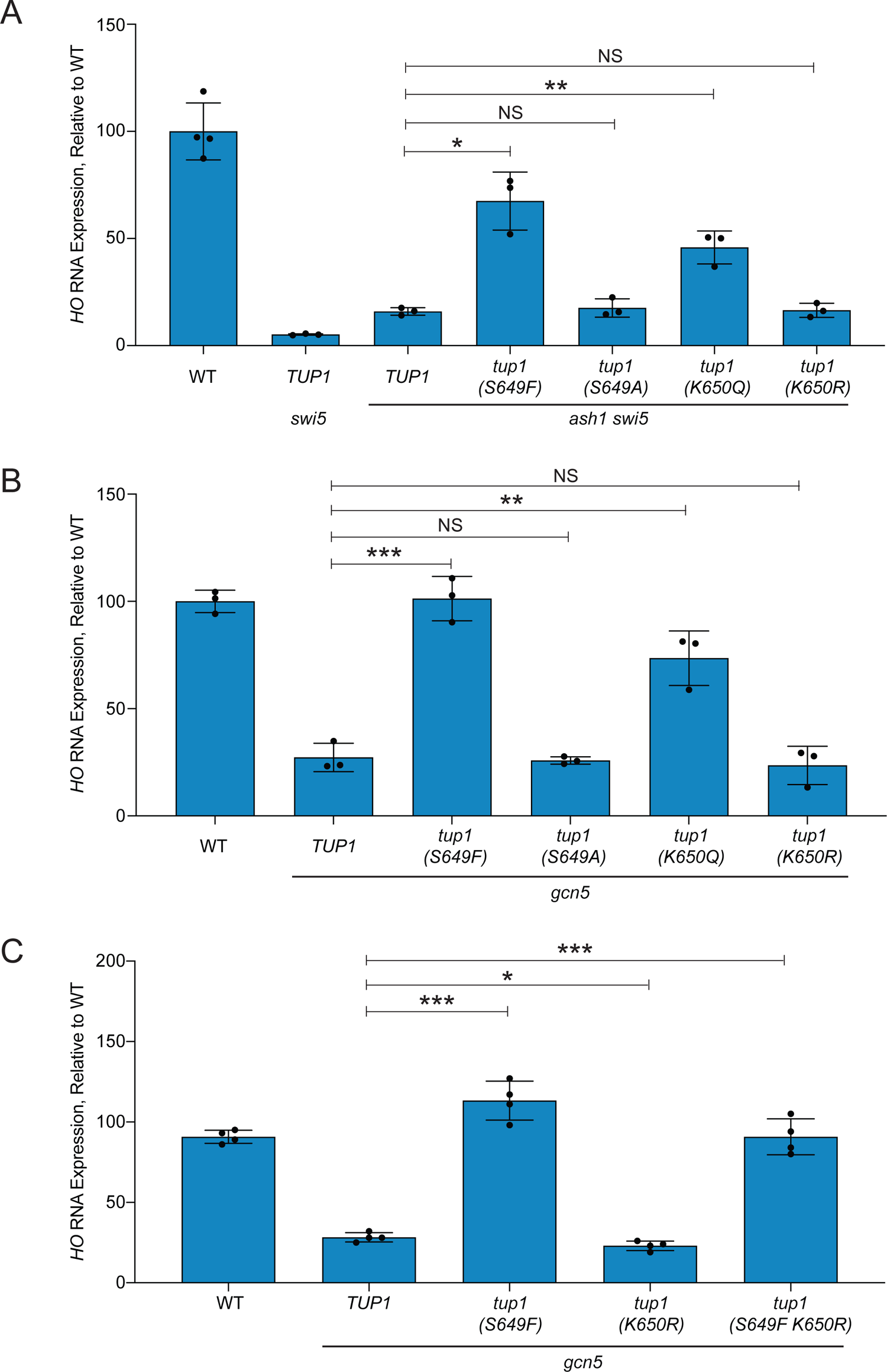
The transcriptional effects of *tup1(S649F)* are not due to changes in post-translational modifications. *HO* mRNA levels were measured by RT-qPCR, normalized to *RPR1*, and expressed relative to wild-type. Each dot represents a single data point, and error bars reflect the standard deviation. *** *p* < 0.001, ** *p* < 0.01, * *p* < 0.05, NS = not significant. (A) The *tup1(S649F)* and *tup1(K650Q)* mutants increase *HO* expression in a *swi5 ash1* strain, while *tup1(S649A)* and *tup1(K650R)* fail to do so. (B) Suppression of *gcn5* is specific to the *tup1(S649F)* and *tup1(K650Q)* alleles. (C) The *tup1(S649F K650R)* double mutant retains the *gcn5* suppression exhibited by the *tup1(S649F)* single mutant.

Two mutations, *tup1(K650Q)* and *tup1(K650R)*, were constructed to mimic either permanent acetylation or deacetylation of K650, respectively. Interestingly, the *tup1(K650Q)* mutant displayed suppression of *HO* expression in the *swi5 ash1* and *gcn5* strains, albeit at a lower level than *tup1(S649F)*, while the *tup1(K650R)* mutant had no effect (Fig 4A, 4B). The difference in expression effects upon substitution of lysine with glutamine or arginine suggests the possibility that acetylation of K650 could play a role in Tup1 regulation. If acetylation of Tup1 were necessary to “turn off” its repressive capacity, then a glutamine permanent acetylation mimic (Q) would be expected to be a weaker repressor. *HO* expression would then increase specifically in the *tup1(K650Q)* mutant, as observed. An arginine permanent deacetylation mimic (R) should not have the same effect.

One hypothesis for the transcriptional effects of *tup1(S649F)* is that it may position K650 such that it can be acetylated but not deacetylated. Tup1 would thus become locked in a “repressor off” state, explaining the similarity between the *tup1(S649F)* and *tup1(K650Q)* mutants. If this scenario explains the phenotype of *tup1(S649F)*, then combination of *tup1(S649F)* with *tup1(K650R)*, which cannot be acetylated, should not allow the increases in *HO* transcription observed in the *tup1(K650Q)* mutant. However, a *tup1(S649F K650R)* double mutant still displayed suppression of a *gcn5* mutant for *HO* expression, suggesting the addition of the bulky phenylalanine group at position 649 has a different effect (Fig 4C). A change in acetylation state is therefore unlikely to explain the transcriptional effects of the *tup1(S649F)* and *tup1(K650Q)* mutants. A more plausible explanation, as suggested above, is that mutation of S649F diminishes Tup1 protein stability by disrupting the 6^th^ WD repeat, reducing the amount of Tup1 available for affecting gene expression. Alternatively, or in addition, the S649F substitution could affect Tup1 interaction with other proteins.

### Many genes are upregulated in *tup1(S649F)*, both in a and α strains

The diminished Tup1(S649F) protein level relative to wild-type suggests this mutant is a hypomorph, expected to display similar but reduced phenotypes and transcriptional changes as a *tup1* null mutant. The slow growth, mating, and flocculation phenotypes of *tup1(S649F)* strains fit this prediction. However, the difference in growth phenotype between **a** and α *tup1(S649F)* mutants has not, to our knowledge, been noted in *tup1* null mutants and suggests expression of Tup1-regulated genes in *tup1(S649F)* may not be identical to that of a *tup1* null. To investigate the nature of transcriptional changes in the *tup1(S649F)* mutants, we performed RNA-Seq analysis, comparing genome-wide expression in wild-type and *tup1(S649F)* strains of both mating types (Table S3). As expected for impaired function of a corepressor protein, both *tup1(S649F)* mating types displayed upregulation of a large number of genes (Fig 5A, Table S4). The α strain had more changes than the **a** strain (561 vs. 359 upregulated genes). Only a small minority of genes in both mating types were downregulated (<10% of the total changed genes), fewer than have been previously observed in a comparison of a *tup1* null with wild-type (Chen *et al*. 2013).

**Figure 5.**
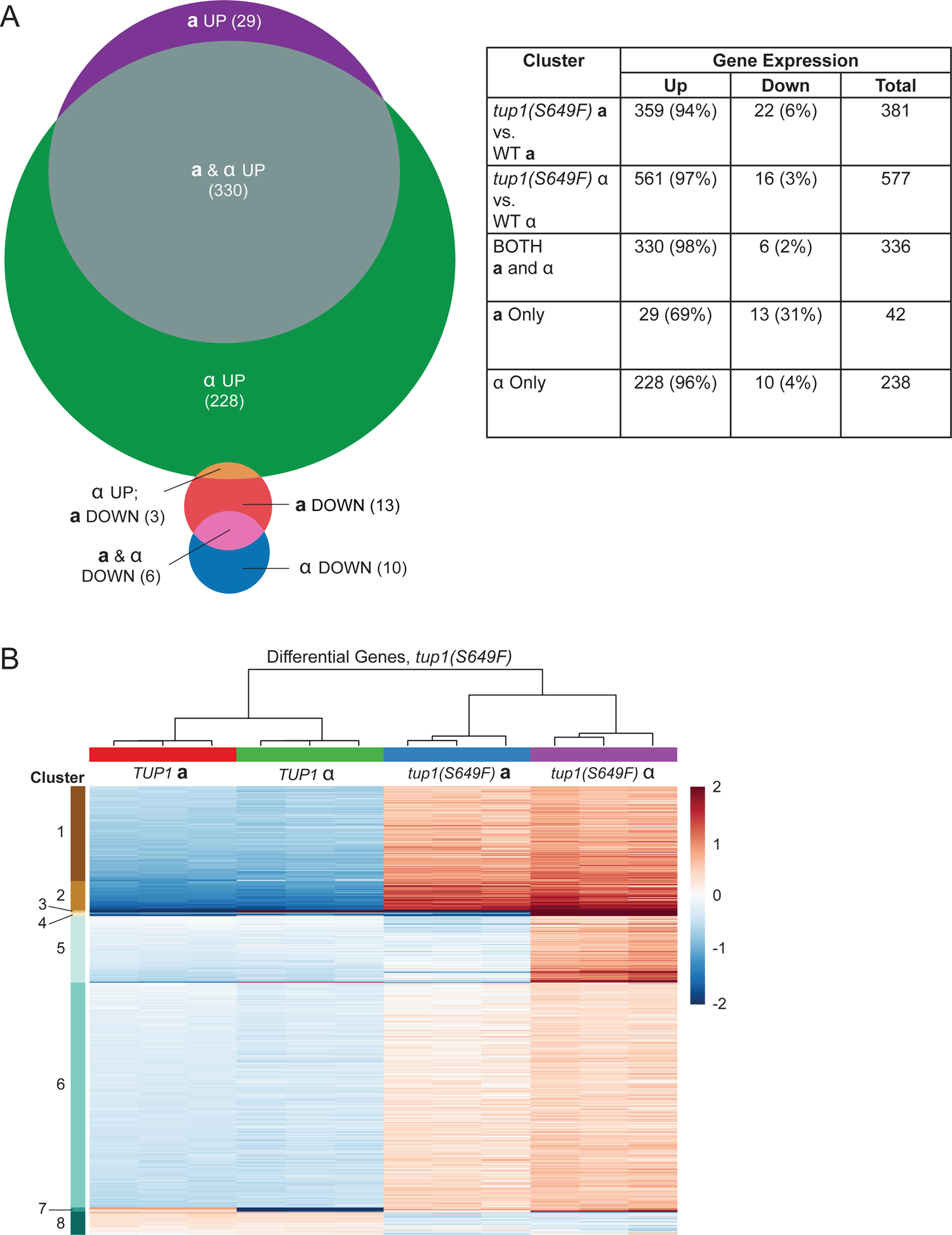
The *tup1(S649F)* mutant affects expression of many genes. (A) A large number of genes show altered expression in both mating types of the *tup1(S649F)* mutant. Protein-coding genes for which RNA-Seq showed a >2-fold increase (“Up”) or decrease (“Down”) in expression were counted for each comparison of *tup1(S649F)* to wild-type (WT) in each mating type. Percentages indicate the proportion of increased or decreased genes out of each horizontal total. Genes indicated as “Both” showed altered expression in both mating types; those indicated as “**a** Only” or “α Only” were mating-type specific. Overlap is shown visually in the Venn diagram at the left. Three genes were not counted in “Both” or “**a**/α Only” due to increased expression in one mating type and decreased expression in the other mating type. (B) k-means clustering highlights different groups of genes altered in the *tup1(S649F)* mutant. The heat map depicts RNA expression levels among the different genotypes listed at the top, expressed as mean centered relative log values, with each row representing an individual gene. Triplicate biological samples for each genotype are shown as individual columns. Only genes that displayed a greater than 2-fold difference in at least one comparison between mutant and wild-type and between **a** and α cells are displayed (642 total). Genes were sorted by an unsupervised k-means clustering algorithm into eight clusters. The hierarchical tree above the heat map indicates the degree of relationship between samples, and the color scale at the right indicates the level of log_2_-fold enrichment.

A heat map showing relative expression values for all differentially expressed genes (*p* < 0.05) that exhibited a greater than 2-fold increase or decrease between mutant and wild-type is shown in Fig 5B. The genes were organized into clusters by a k-means algorithm, using eight clusters to emphasize the pattern of gene expression changes between strains. Clustering showed groups of genes with increased expression in both mating types in the *tup1(S649F)* mutant strains (Clusters 1, 2, and 6) as well as one group with decreased expression (Cluster 8). Cluster 5 highlighted genes for which expression increased only in the α mating type of the mutant strain. The small number of genes that were differentially expressed in **a** vs. α strains, either in wild-type or mutant or both, comprised Clusters 3, 4 and 7 (discussed below). A comprehensive list of all genes with greater than a 2-fold change in expression in mutant vs. wild-type for both mating types is provided in Table S4.

Gene ontology (GO) analysis identified several categories of genes that displayed increased expression in *tup1(S649F)* in each mating type (Table S5). Consistent with previous reports, these included genes involved in carbohydrate metabolism, oxidation/reduction, transmembrane transport and cell wall organization (Derisi *et al*. 1997; Green and Johnson 2004). In addition, transcription of some DNA-binding transcription factors was altered. A large number of genes that were altered in the *tup1(S649F)* mutant encoded proteins found at the cell periphery. Genes shown previously to be regulated by Tup1 via different repressor proteins were found to be upregulated, including **a**-specific *MFA2*, glucose-repressed *SUC2*, oxygen-repressed *ANB1*, *HEM13*, and *DAN1*, glucose-regulated *HXT* genes, DNA damage response genes *RNR2*, *3*, *4,* and flocculation *FLO* genes. The effect of *tup1(S649F)* on these diverse pathways suggests a general loss of repression by the mutant protein. All of the same categories of genes were identified when comparing *tup1(S649F)* α expression to wild-type α expression. However, mutant α strains exhibited additional changes in genes involved in small molecule and organic acid metabolic processes, cellular response to chemical stimuli, and sexual reproduction (Table S5). As suspected from the differential growth phenotype in *tup1(S649F)* **a** and α cells relative to *tup1* null, many changes in the *tup1(S649F)* α strain relative to wild-type were not identified in a previous comparison of *tup1* null and wild-type α strains (discussed further below; Chen *et al*. 2013). Performing GO analysis on individual k-means clusters from Fig 5B failed to identify additional categories of genes regulated by Tup1 or to demonstrate that particular clusters contained a high percentage of genes with similar functions.

### Tup1(S649F) shows increased association at many genomic locations

The reduced level of Tup1(S649F) protein relative to wild-type, particularly in the α strain (Fig S1), suggests the transcriptional effects of *tup1(S649F)* could be due to less Tup1 protein bound within target promoter regions. General loss of Tup1 protein across the genome could explain the observed upregulation of Tup1 target genes controlled by different repressors and would suggest *tup1(S649F)* largely behaves similar to a null mutant. This is supported by the observation that the *tup1(S649F)* mutation reduced but did not eliminate Tup1 occupancy in a ChIP assay to its most predominant site in the *HO* promoter (Fig 6A, Top). However, association of Tup1(S649F) with two other target promoters, *POG1* and *TEC1*, was increased relative to wild-type (Fig 6A, Middle & Bottom). The increased occupancy of Tup1(S649F) is difficult to reconcile with the observations of a reduced level of protein relative to wild-type and the upregulation of many genes. We therefore performed ChIP-Seq to investigate the association of Tup1 and Tup1(S649F) with their genome-wide targets in both **a** and α mating types.

**Figure 6.**
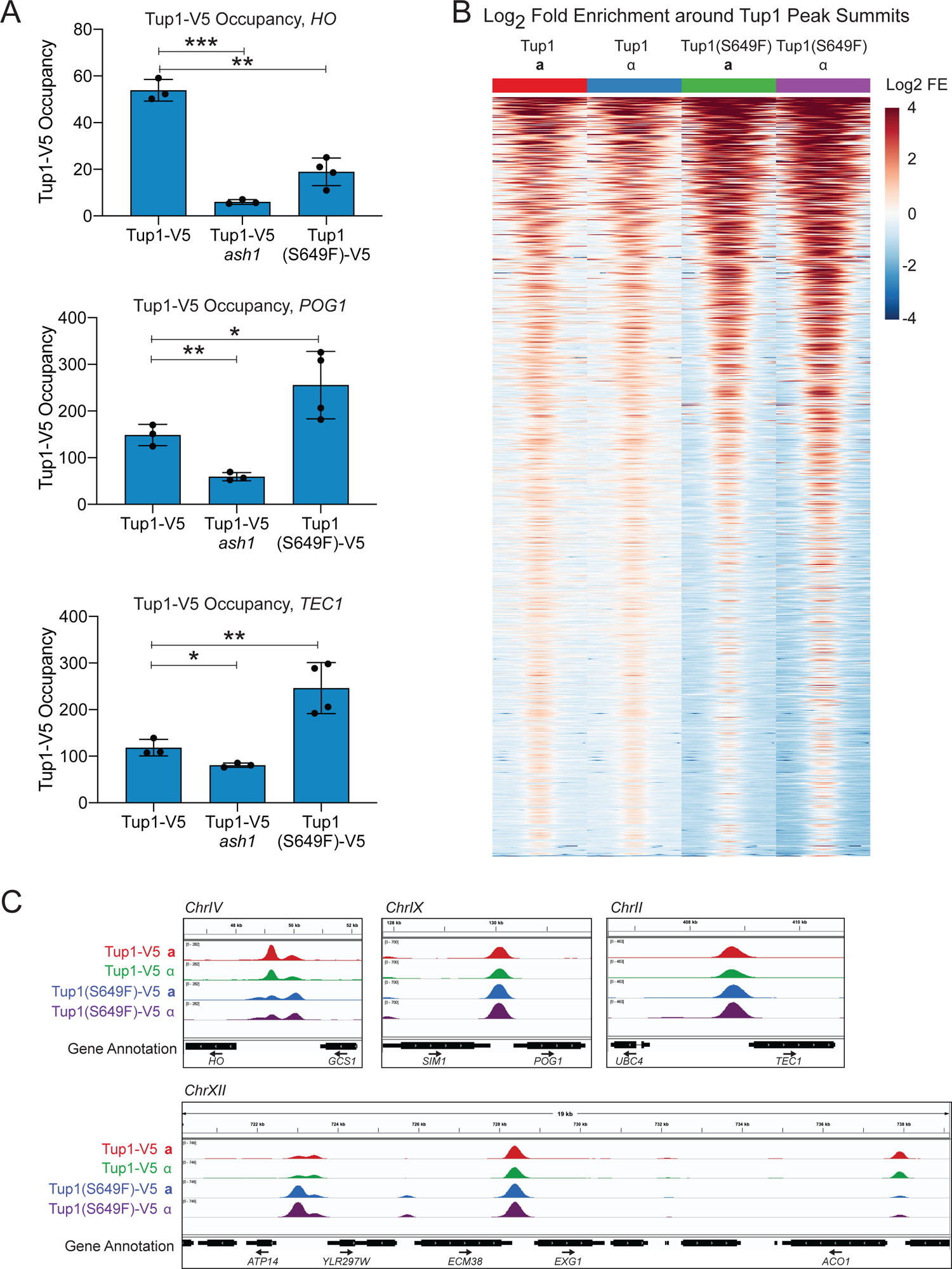
The *tup1(S649F)* mutant displays increased occupancy relative to wild-type at many genomic locations. (A) Tup1(S649F) occupancy decreased at the *HO* promoter but increased at the intergenic regions upstream of *POG1* and *TEC1*. Tup1-V5 ChIP analysis, showing enrichment upstream of the *HO* (−1295 to −1121), *POG1* (−570 to −435) and *TEC1* (−433 to −259) genes. For each sample, occupancy was normalized to its corresponding input DNA and to a No Tag control. Each dot represents a single data point, and error bars reflect the standard deviation. *** *p* < 0.001, ** *p* < 0.01, * *p* < 0.05. (B) Occupancy of Tup1(S649F) appears to be higher than wild-type Tup1 occupancy at most genomic locations. Heat maps depict the log_2_-fold enrichment of Tup1-V5 and Tup1(S649F)-V5 at peak summits genome-wide (1266 peaks), displaying enrichment from −625 to +625 relative to the center of each Tup1-V5 peak, in bins of 25-bp. The data for each strain represents an average of three biological replicates, and the color scale at the right indicates the level of log_2_-fold enrichment. Each horizontal line depicts a single Tup1-V5 peak. (C) Tup1(S649F) occupancy shows variation among individual genomic sites. Snapshots of ChIP-Seq results from the Genome Browser IGV (Broad Institute) show the sequenced fragment pileups for the intergenic regions upstream of *HO* (Top Left), *POG1* (Top Middle) and *TEC1* (Top Right), and a region of chromosome XII (Bottom). The colored tracks show ChIP-Seq results for Tup1-V5 WT **a** (red), Tup1-V5 WT α (green), Tup1(S649F)-V5 **a** (blue) and Tup1(S649F)-V5 α (purple). The bottom track displays gene annotations. The four samples were group autoscaled independently at each location, to account for differences in enrichment between sites. Tracks represent an average of three biological replicates.

Distinct differences were observed between ChIP-Seq samples of Tup1 wild-type and Tup1(S649F) of both mating types conducted with triplicate biological samples, as judged by Pearson correlation (Fig S2). In contrast to the expectation that Tup1 occupancy would be reduced in the Tup1(S649F) mutant, we observed an overall stronger association of the mutant Tup1 protein with chromatin than wild-type Tup1, visualized by higher log_2_ fold enrichment at peaks across the genome (Fig 6B). This suggests that the *POG1* and *TEC1* upstream regions tested are reflective of the genome-wide changes in Tup1 occupancy in the *tup1(S649F)* mutant. At the *HO* promoter, ChIP-Seq showed that while the major peak of Tup1 occupancy at *HO* was reduced in *tup1(S649F)*, as we had observed in the qPCR analysis (Fig 6A), the minor peak of Tup1 occupancy at *HO* actually increased in the mutant strain, similar to those within the *POG1* and *TEC1* upstream regions (Fig 6C, ChrIV).

Visualization of ChIP-Seq fragment densities across the genome with the IGV genome browser (Broad Institute) suggested that Tup1 occupancy was redistributed in the *tup1(S649F)* mutant relative to wild-type. While many locations displayed increased occupancy in the mutant relative to wild-type, some appeared mostly unaltered, and some had decreased occupancy (Fig 6C). New peaks seemed to appear in the mutant, but careful observation in a genome browser showed that these locations had a small amount of Tup1 occupancy in wild-type that became much more predominant in the *tup1(S649F)* mutant (Fig S3A). These genomic results demonstrate that decreased Tup1 protein stability did not cause a simple loss of occupancy uniformly across the genome, but paradoxically resulted in many regions of increased association. Thus, the *tup1(S649F)* mutant is clearly distinct from a *tup1* null.

The majority of Tup1 peaks in both wild-type and mutant were located within intergenic regions, as expected (770/1266, 60%; Fig S3B). Approximately 24% of peaks were located predominantly over untranslated regions (UTRs) and/or open reading frames (ORFs). Another 16% overlapped tRNA and other Pol III genes (*RPR1*, *SCR1*) and snRNAs. Peaks in intergenic regions displayed a range of log_2_-fold changes in occupancy between Tup1(S649F) strains and wild-type Tup1 strains, whereas most ORF and Pol III / snRNA locations showed decreased occupancy in the mutant relative to wild-type, even more so in α strains than in **a** strains (Fig S3C, S3D). Occupancy over tRNA and snRNA genes was weak relative to the majority of Tup1 peaks; therefore, it is uncertain what role Tup1 plays at these loci (Fig S3C). However, with over 50% of yeast tRNA genes showing occupancy of Tup1 within our threshold parameters and a number of others with weaker but observable association in a genome browser, the occupancy of Tup1 peaks at Pol III genes is unlikely to be coincidental.

### Many genes with altered expression in the *tup1(S649F)* mutant are located downstream of a Tup1 peak

Current models of Tup1 function suggest it affects transcription by associating with target promoters and influencing their chromatin state and recruitment of factors. Thus, Tup1 occupancy is expected within intergenic regions upstream of the genes that its complex directly regulates. To assess how many of the genes with altered expression in the *tup1(S649F)* mutant could be regulated by Tup1 directly, we intersected the list of differentially expressed genes with those that had a Tup1 peak located upstream of the ORF (Table 2). tRNA genes were excluded from this analysis due to their small size and inability to accurately map sequencing reads to specific loci. In both mating types, the majority of protein-coding genes that displayed increased expression in the mutant were located downstream of a Tup1 peak (78% for **a**, 67% for α). Decreased expression changes appeared more likely to be indirect effects, at least in **a** strains, as the few genes with diminished expression in the mutant strains were not as likely to be associated with Tup1 occupancy (29% in **a**; 56% in α). Overall, 75% of genes with altered expression in the *tup1(S649F)* mutant in **a** strains and 66% in α strains had an upstream Tup1 peak, suggesting most transcriptional effects are a result of changes to the Tup1 located within their promoters. A small number of genes with expression changes in both mating types had a Tup1 peak located over the ORF; based on current models of Tup1 function, it is not clear if or how Tup1 could be regulating these genes directly.

**Table 2.** Location of Tup1 peaks relative to genes that change >2-fold in the *tup1(S649F)* mutant relative to WT.

A. Strains with mating type **a**.
| Location of Tup1 Peak Relative to Gene<sup>a</sup> | No. of Genes that Increase Expression | No. of Genes that Decrease Expression | Total Genes with Expression Changes |
| --- | --- | --- | --- |
| Upstream<sup>b</sup> | 280 (78%) | 6 (29%) | 286 (75%) |
| Over the ORF | 3 (1%) | 4 (19%) | 7 (2%) |
| No Peak<sup>c</sup> | 77 (21%) | 11 (52%) | 88 (23%) |
| Total | 360 (100%) | 21 (100%) | 381 (100%) |

B. Strains with mating type $\alpha$ .
| Location of Tup1 Peak Relative to Gene<sup>a</sup> | No. of Genes that Increase Expression | No. of Genes that Decrease Expression | Total Genes with Expression Change |
| --- | --- | --- | --- |
| Upstream<sup>b</sup> | 374 (67%) | 9 (56%) | 383 (66%) |
| Over the ORF | 12 (2%) | 1 (6%) | 13 (2%) |
| No Peak<sup>c</sup> | 175 (31%) | 6 (38%) | 181 (31%) |
| Total | 561 (100%) | 16 (100%) | 577 (100%) |

<sup>a</sup> See Methods for description of how these were determined.
<sup>b</sup> Includes both locations in which a peak was only present upstream of the gene of interest (217 for **a**; 300 for $\alpha$ ), as well as those in which peaks flanked the gene of interest (63 for **a**; 74 for $\alpha$ ).
<sup>c</sup> Includes locations in which a peak was observed downstream of the gene of interest.

### Increased association of Tup1(S649F) correlates with increased expression of downstream genes

To examine patterns in occupancy changes between the strains, all of the Tup1 peaks were merged into a single set and a mean enrichment score obtained for each strain, followed by k-means clustering of the data. This analysis identified groups of Tup1 peak locations that displayed differences in occupancy between the wild-type and mutant strains. Four of the eight clusters in Fig 7A showed increased occupancy of Tup1(S649F) in both **a** and α strains (clusters 2, 4, 5, and 6), while two other clusters had diminished occupancy of the Tup1(S649F) relative to WT (clusters 3 and 8). The remaining two clusters appeared to have only minor or variable occupancy changes in Tup1(S649F) relative to wild-type (clusters 1 and 7). Intergenic peaks were disproportionally represented in the clusters in which occupancy appeared to increase in mutant relative to wild-type (greater than 80% of the peaks; clusters 2,4,5 and 6; Table S6). However, those clusters with diminished occupancy in mutant relative to wild-type had many fewer intergenic peaks (23% for cluster 3; 35% for cluster 8). Thus, locations that lost occupancy in the mutant tended to be those with occupancy in non-intergenic locations, whereas those that gained occupancy in the mutant tended to be within intergenic positions from which they could influence expression of downstream target genes.

**Figure 7.**
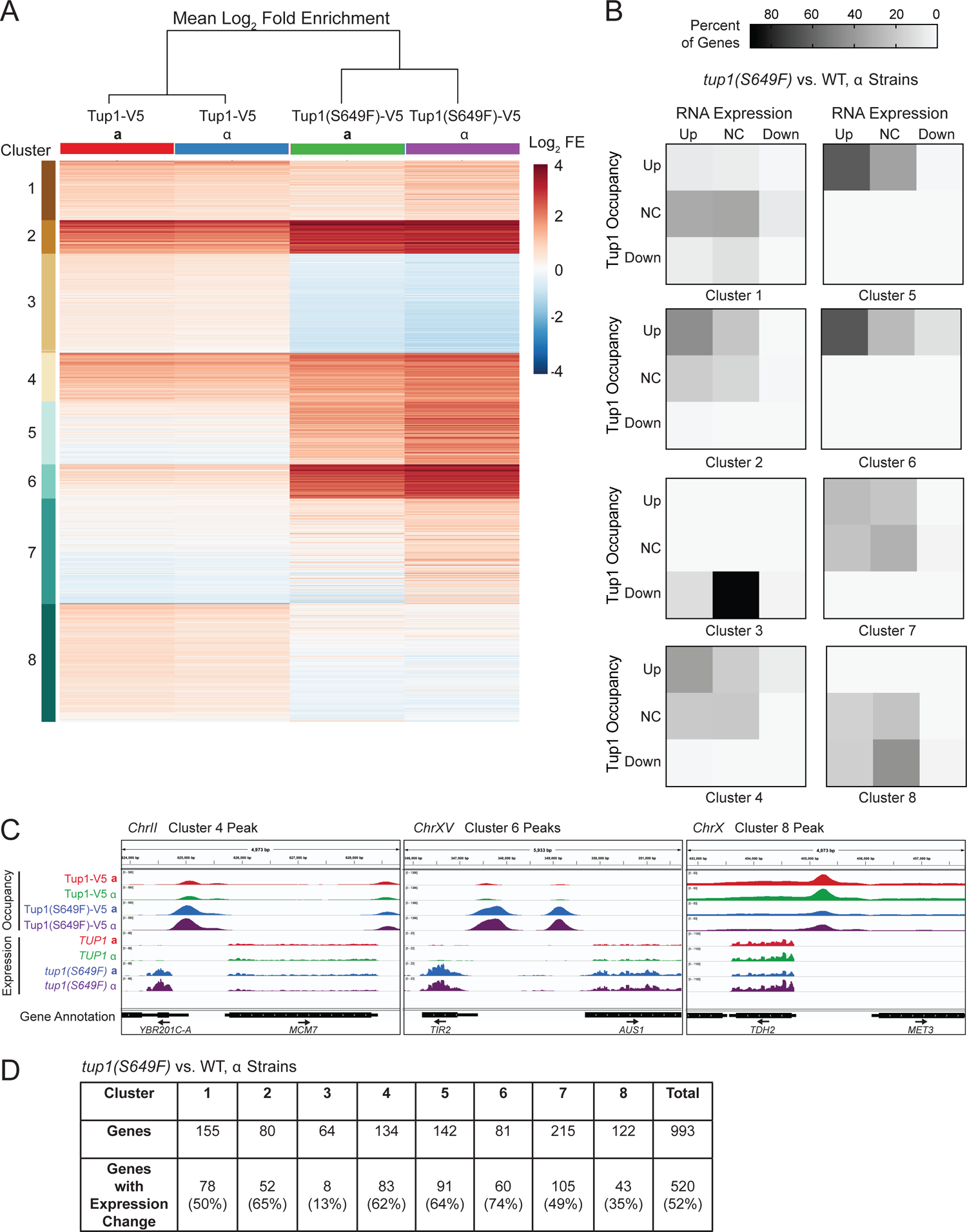
Gene expression changes tend to occur at Tup1 intergenic sites with increased Tup1(S649F) occupancy. (A) Groups of Tup1 peaks show differences in Tup1 vs. Tup1(S649F) occupancy. The heat map depicts the mean Tup1 log_2_-fold enrichment among the different genotypes listed at the top (each an average of triplicate biological samples). The rows represent sites of Tup1 occupancy (1266 total). Tup1 peaks were sorted by an unsupervised k-means clustering algorithm into eight clusters. The hierarchical tree above the heat map indicates the degree of relationship between samples. (B) Increased Tup1-S649F occupancy often occurs in conjunction with increased expression of downstream genes. Each square within the 3 x 3 heat maps represents a unique combination of Tup1 occupancy and RNA expression of the downstream gene, where “Up” and “Down” indicate increased or decreased occupancy/expression in *tup1(S649F)* α vs. wild-type α, respectively, and “NC” (No Change) indicates less than 1.5-fold alteration in levels. The color scale at the top indicates the percentage of genes in each category. Genes were separated by cluster from (A) and were only included if they were located downstream of a Tup1 peak and had a valid expression level from RNA-Seq (993 total). Peaks were counted twice if present between divergent genes. Some genes were counted more than once, due to two or three Tup1 peaks within the intergenic region upstream of the gene (53 total). (C) Individual Tup1 occupancy sites show different effects on expression of downstream genes. Snapshots of ChIP-Seq (Occupancy) and RNA-Seq (Expression) results from the Genome Browser IGV (Broad Institute) show the sequenced fragment pileups for representative peaks on chromosomes II, XV and V (examples from clusters 4, 6, and 8). Tracks represent an average of three biological samples for each strain. The four strains for each experiment were autoscaled as a group for occupancy and expression separately to account for differences in fragment depth between locations. Colors for both ChIP-Seq and RNA-Seq are as follows: Tup1-V5 WT **a** (red), Tup1-V5 WT α (green), Tup1(S649F)-V5 **a** (blue) and Tup1(S649F)-V5 α (purple). The bottom track displays gene annotations. Gene names are indicated for those located downstream of a Tup1 peak. (D) Approximately half of genes downstream of Tup1 peaks show altered expression in the *tup1(S649F)* α mutant. Shown are the number and percentage of genes downstream of Tup1 peaks in each cluster from (A) with >1.5-fold change in RNA-Seq values between *tup1(S649F)* α and wild-type α.

We next investigated the relationship between changes in Tup1 association and gene expression changes in *tup1(S649F)* vs. wild-type. For this analysis, we considered only genes that are located downstream of Tup1 peaks and used the ChIP k-means clusters as a method to divide the large number of gene points into smaller groups for easier interpretation, while also highlighting the differences among clusters. Log_2_ fold change in RNA expression was plotted vs. log_2_ fold change in Tup1 occupancy for individual gene/peak combinations (Fig S4, S5). Each gene was then categorized as “Up” or “Down” for both ChIP occupancy and RNA expression if the absolute change was greater than 1.5-fold change, otherwise as “NC” (No Change). The simplified heat maps in Fig 7B allow visualization of which occupancy and expression combinations were most prevalent in each cluster, when comparing mutant and wild-type strains with an α mating type. Each of the nine squares represents a unique combination of “Up,” “NC,” or “Down,” in Tup1 occupancy and in gene expression. Notably, none of the clusters had significant numbers of genes in the upper right square, which would indicate increased Tup1 occupancy and decreased expression, an expectation for a strict corepressor. Instead, increases in occupancy often occurred concomitantly with increased expression. In clusters 2, 4, 5 and 6, the upper left square had a substantial number of genes (genomic snapshot examples of cluster 4 and 6 peaks are in Fig 7C). Clusters 1 and 8, as expected, showed the largest proportion of genes in the central square, representing no significant changes either in occupancy or expression levels (example of cluster 8 peak in Fig 7C). Finally, cluster 3, which exhibited mostly decreases in Tup1 occupancy, had very few changes in gene expression. Again, this does not agree with the prediction for a corepressor, in which diminished association with the promoter would be expected to increase expression of target genes. Similar results were obtained for **a** strains (Fig S6).

Overall, those clusters which exhibited increased occupancy of Tup1 (2, 4, 5, 6) had the largest proportion of downstream genes with significantly altered expression levels, approximately 50-60% (Fig 7D), with the vast majority of those changes representing increased expression (Fig S4, S5). Considering that our analysis was performed in rich media conditions in logarithmically growing cells representing all stages of the cell cycle, this level of positive correlation between Tup1 occupancy and gene expression is likely significant. The actual number of genes displaying changes in occupancy and expression may be higher than the numbers reported here; viewing the genes as individual points revealed that genes within the “No Change” category often displayed minor alterations in occupancy and expression (Fig S4, S5). These results suggest that while the Tup1(S649F) protein is less abundant and less stable than wild-type, it associates strongly with many genic loci with increased expression. Thus, many of the RNA expression changes observed at locations downstream of Tup1 peaks may be a direct result of Tup1(S649F) mutant properties.

### The *tup1(S649F)* α strain fails to down-regulate critical a-specific genes

The mating irregularities and shmoo phenotype of the *tup1(S649F)* α strain suggest mis-regulation of the genes normally expressed in only one of the two haploid mating types. The three *S. cerevisiae* cell types, haploid **a** cells, haploid α cells and **a**/α diploid cells, each have distinct patterns of mating-type specific transcription (Haber 2012).

Genes specific to the **a** mating type are activated in **a** cells by Mcm1 (Elble and Tye 1991). In α cells and **a**/α diploids, Mcm1 heterodimerizes with α2 to form a repressor that turns off expression of these **a**-specific genes (Johnson and Herskowitz 1985; Keleher *et al*. 1988). Haploid-specific genes, such as *HO*, are repressed in diploids by a heterodimeric **a**1/α2 repressor (Goutte and Johnson 1988; Keleher *et al*. 1988). Repression by both Mcm1/α2 and **a**1/α2 is dependent on Tup1 (Keleher *et al*. 1992). An altered function of Tup1 in haploid cells could therefore lead to aberrant expression of **a**-specific genes in the α cells.

To investigate this prediction, we first used the RNA-Seq analysis to identify genes that were differentially expressed between **a** and α haploid cells. Only thirteen genes showed an expression change greater than 2-fold between the mating types in wild-type. These genes are listed in Table S8 and include those encoding the **a** and α mating pheromones [*MFA1*, *MFA2*, *MF(ALPHA)1*, *MF(ALPHA)2*], the receptors for **a**- and α-factor (*STE3*, *STE2*), and other genes encoding proteins involved in the pheromone response. The volcano plots in Fig 8A, which display significance vs. fold change, highlight the changes in these thirteen genes between the two mating types (noted with gene common names). Genes noted on the left side of the graph are **a**-specific genes that were down-regulated in α cells (*MFA1*, *MFA2*, *BAR1*, *STE2*, *STE6* and *AGA2*), and genes noted on the right side of the graph are α-specific genes that were expressed only in α cells [*MF(ALPHA)1*, *MF(ALPHA)2*, *STE3*, *AFB1*, *SAG1*, *PRM7*, *BSC1*]. These results are consistent with prior studies that have examined differential expression between **a** and α cells, with the addition of *BSC1* and *PRM7* that have smaller transcriptional effects (but still above the 2-fold threshold; Galgoczy *et al*. 2004). *BSC1* and *PRM7* should be considered as a single gene for our analyses, as the two comprise a continuous ORF, *IMI1*, in W303 strains (but they are separate genes in the S288C reference strain and are listed separately in Tables S3, S4, and S7).

**Figure 8.**
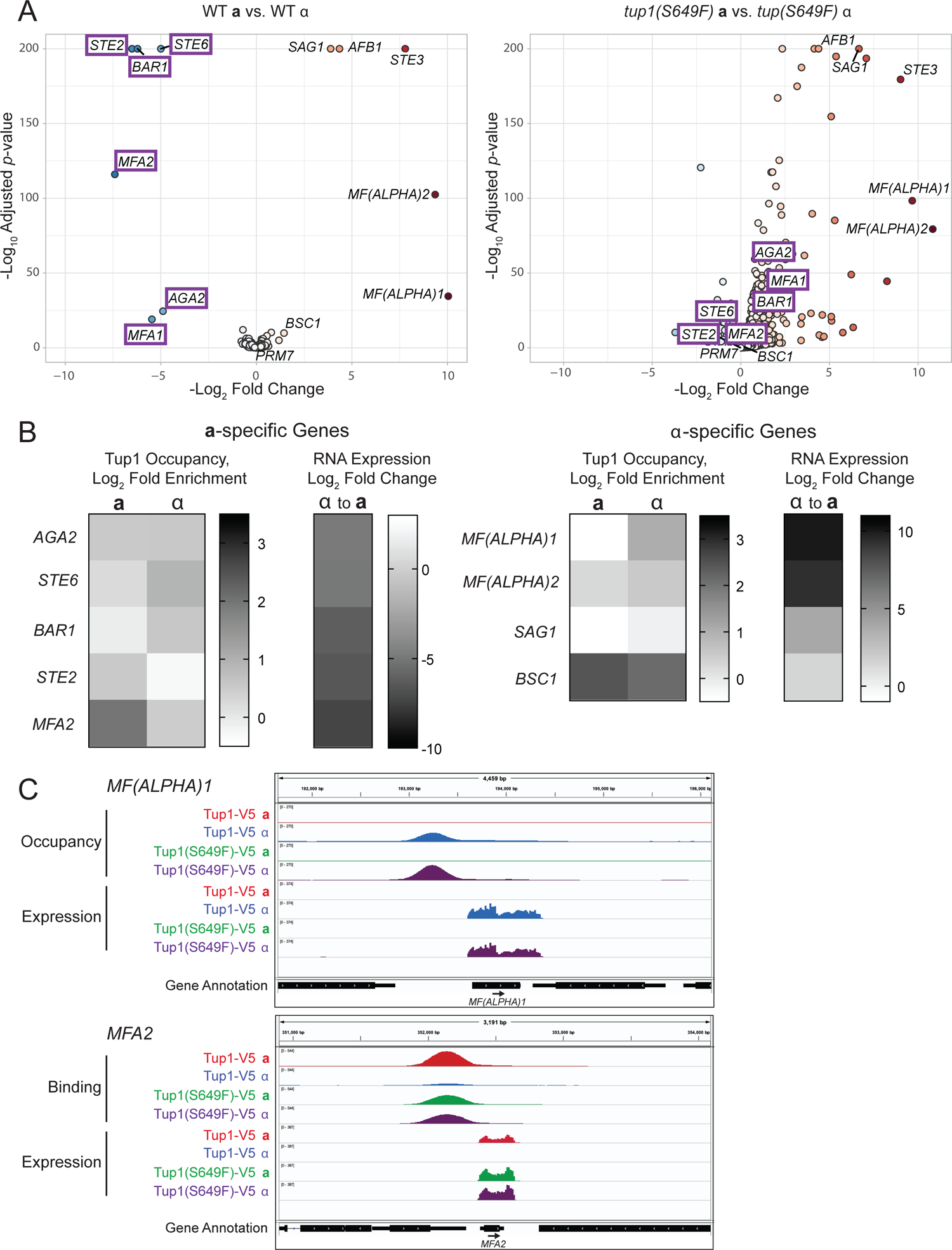
Regulation of mating type genes is altered in the *tup1(S649F)* mutant. (A) Many more genes show differential regulation between **a** and α mating types in the *tup1(S649F)* mutant than in wild-type. Volcano plots of RNA-Seq data show expression differences between **a** and α strains for wild-type (Left) and the *tup1(S649F)* mutant (Right). The -log_10_ adjusted *p* values were plotted relative to the log_2_ fold change for all genes (n = 5730). Each dot represents a single gene; those with increased or decreased expression are shown as deepening shades of red or blue, respectively. The thirteen genes identified with their common name are differentially regulated greater than 2-fold between the wild-type **a** and α strains. The six **a**-specific genes that lose their differential expression between mating types in the *tup1(S649F)* mutant are boxed in purple. (B) Tup1 occupancy at mating-type specific genes does not predict RNA expression changes. Shown are heat maps indicating log_2_-fold enrichment of Tup1 occupancy and log_2_-fold change in RNA expression at **a**-(Left) and α-specific genes (Right). Color scales are shown to the right of each heat map. Only genes with detectable ChIP-Seq peaks in wild-type **a** and/or α strains are shown. Genome browser snapshots for all genes are shown in Fig S7, and numerical values are listed in Table S8. (C) *MF(ALPHA)1* and *MFA2* unexpectedly display upstream Tup1 occupancy in the mating type strain in which the gene is expressed rather than repressed. Snapshots of ChIP-Seq (Occupancy) and RNA-Seq (Expression) results from the IGV genome browser (Broad Institute) show the sequenced fragment pileups for Tup1 peaks in the intergenic regions upstream of *MF(ALPHA)1* and *MFA2*. Tracks represent an average of three biological samples for each strain. The four strains for each experiment were autoscaled as a group for occupancy and expression separately to account for differences in fragment depth between locations. Colors for both ChIP-Seq and RNA-Seq are as follows: Tup1-V5 WT **a** (red), Tup1-V5 WT α (blue), Tup1(S649F)-V5 **a** (green) and Tup1(S649F)-V5 α (purple). The bottom track displays gene annotations.

The plot for *tup1(S649F)* α vs. **a** strains is strikingly different and illustrates that many more genes showed differential expression between the mating types in the mutant relative to wild-type (Fig 8A, Right). A large number of genes appeared on the right side of the graph, indicating increased expression in α cells relative to **a** cells. A smaller number of genes appeared on the left side of the graph; these genes that showed diminished expression in α cells did not have changes that were as dramatic as those that increased on the right side. The thirteen genes with differential expression in wild-type **a** and α cells are also indicated with their common names on the *tup1(S649F)* plot. Consistent with the expectation that Tup1 represses expression of **a**-specific genes, the six **a**-specific genes that are normally substantially downregulated in α cells were either less downregulated (*STE2*, *STE6*, *MFA2*) or were now upregulated (*BAR1*, *MFA1*, *AGA2*) in *tup1(S649F)* α cells (Bold purple boxed genes in Fig 8A; Fold change α vs. **a** for wild-type and mutant listed in Table S8). Only 20% of the genes we found to be aberrantly differentially expressed in *tup1(S649F)* α also displayed changes in gene expression in a *tup1* null α strain (Chen *et al*. 2013).

Gene ontology analysis of genes with altered expression in *tup1(S649F)* α vs. *tup1(S649F)* **a** identified most of the same categories that were previously found in the comparison of *tup1(S649F)* α to wild-type α that were unique to α strains (Table S5). The percentage of genes involved in aspects of mating, including conjugation with cellular fusion and response to pheromone, increased in the *tup1(S649F)* α vs. *tup1(S649F)* **a** pool relative to the comparison to wild-type α and therefore had much more significant *p*-values; this was not true of the other α-specific gene sets. Manual examination and curation of the genes that became differentially regulated between α and **a** strains in the *tup1(S649F)* mutant identified a few additional genes involved in various aspects of mating, for 18 total genes (in addition to the 13 that change in WT **a** vs. α cells, total = 31; Table S8). Thus, the pool of genes regulated by Tup1 that are involved in mating appears to extend beyond the set of traditionally defined **a**-specific genes. The promoters of these additional genes are distinct from the **a**-specific genes because they are not bound by α2 (Galgoczy *et al*. 2004). All but two of them did not have Tup1 occupancy above our threshold of detection in wild-type **a** or α strains.

However, five more had detectable Tup1(S649F) occupancy in the α strain. While expression changes for these few genes could be directly due to Tup1(S649F) occupancy within their promoters, the majority of changes appear to be indirect effects. Nevertheless, aberrant expression of these additional genes may contribute to the α-specific mating irregularities and growth defect observed in the *tup1(S649F)* mutant.

### Wild-type Tup1 binds upstream of some expressed mating-type specific genes

We expected the wild-type Tup1 ChIP-Seq to show occupancy of Tup1 upstream of **a**-specific genes in the α strain, as Tup1 is required for repression of these genes via Mcm1/α2 (Keleher *et al*. 1992). However, only two promoters, those of *STE6* and *BAR1*, behaved as predicted, with stronger Tup1 occupancy in the α strain than in the **a** strain (Fig 8B, S7). *MFA2* and *STE2* displayed the opposite pattern and had a larger amount of upstream Tup1 in the **a** strain in which the gene is expressed rather than repressed. The final two genes, *AGA2* and *MFA1*, had similar occupancy in **a** and α strains (*MFA1* is not shown in Fig 8B because it fell below the threshold detection of our peak calling program, but very weak occupancy could be seen visually in a genome browser; Fig S7). Tup1 occupancy within the promoters of the **a**-specific genes was generally weak, with the exception of occupancy upstream of *MFA2* in the **a** cell, and failed to correlate with the dramatic difference in RNA expression observed for these genes between the **a** and α strains (Fig 8B, S7; Table S8). Thus, Tup1 did not show the differential association at **a**-specific promoters between mating types predicted by the long-standing model that Tup1-Cyc8 is recruited to these promoters in α cells for repression via α2.

The α-specific genes were not necessarily expected to display upstream Tup1 occupancy, since their differential expression between **a** and α strains is driven by the presence of the α1 activator specifically in α cells rather than by repression in **a** cells (Strathern *et al*. 1981). However, *MF(ALPHA)1*, *MF(ALPHA)2*, *SAG1* and *BSC1/PRM7* all showed upstream Tup1 occupancy in at least one strain (Fig 8B, S7). The Tup1 signal upstream of *BSC1/PRM7* appears as 2-3 peaks and is the strongest of all **a**- and α-specific genes. The entire group of mating type-specific genes, both **a**- and α-specific, therefore shows a range of Tup1 occupancy within their promoters, rather than the expected Tup1 association upstream of only the **a**-specific genes in α cells.

Most intriguing among the mating-type specific genes were the **a**-specific gene *MFA2* and the α-specific gene *MF(ALPHA)1*. Both of these promoters exhibited strong Tup1 occupancy in the mating type in which the gene is strongly expressed [**a** for *MFA2* and α for *MF(ALPHA)1*; Fig 8C]. *MFA2* also displayed Tup1 occupancy in the α cell, as expected, but the association was weaker than in the **a** cell. In the *tup1(S649F)* mutant, the *MFA2* promoter showed strong occupancy of Tup1 in both **a** and α cells and a loss of differential RNA expression between the two mating types. Tup1 occupancy and RNA expression of α-specific genes *BSC1/PRM7*, *SAG1*, and *MF(ALPHA)2* all increased in the α strain in the *tup1(S649F)* mutant (Fig S7; Table S8). Collectively, these results suggest Tup1 and Tup1(S649F) could act as a coactivator at some of the mating-type genes. The critical distinction is that for *MFA2* and *MF(ALPHA)1*, we observed this coactivator function as a feature of wild-type differential expression between mating types, suggesting that even in a wild-type cell, Tup1 shows properties consistent with coactivator function. These results are difficult to reconcile with current models of Tup1 repression of **a**/α regulation and suggest the role of Tup1 at these genes may be different than previously hypothesized.

## DISCUSSION

The Tup1-Cyc8 complex was originally defined as a corepressor due to its ability to decrease transcription when tethered upstream of reporter genes. However, increasing evidence shows that Tup1 and Cyc8 display properties of coactivators in certain contexts, suggesting the complex would be more aptly termed a coregulator. We present here two additional observations that contribute to the body of evidence indicating coactivator function of Tup1-Cyc8. First, increased occupancy of the Tup1(S649F) mutant protein is correlated with upregulation of downstream target genes. Second, occupancy of wild-type Tup1 upstream of two mating-type specific genes, *MFA2* and *MF(ALPHA)1*, occurs within the strain in which the gene is expressed rather than in the strain in which the gene is repressed. These points necessitate re-evaluation of models of how Tup1 functions and the mechanisms responsible for mating type specific gene expression.

### The *tup1(S649F)* mutant appears to favor the coactivator state

When considering simply the RNA-Seq results, *tup1(S649F)* behaves as expected for a corepressor mutation, as most genes show increased expression relative to wild-type (Fig 5). However, the addition of Tup1 occupancy data from ChIP-Seq presents a more complex picture, in which increased gene expression is observed even at locations in which the mutant protein is more abundant than the wild-type protein (Fig 7). We suggest that at some promoters, the *tup1(S649F)* mutant not only has reduced ability to perform its corepressor function but has shifted toward its coactivator state. When compared to wild-type, many fewer genes exhibit decreased expression in a *tup1(S649F)* mutant than in a *tup1* null, a distinction that is also consistent with retention of a coactivator function by *tup1(S649F)* (Chen *et al*. 2013).

The preference of Tup1(S649F) for the coactivator state could be explained by reduction or loss of its association with DNA-binding recruiter proteins at many target promoters. Several studies point to disruption of this interaction between Tup1-Cyc8 and its DNA-binding partner as a mechanism by which the complex is converted from a corepressor to a coactivator, often via phosphorylation of the DNA-binding protein (Papamichos-Chronakis *et al*. 2002; Proft and Struhl 2002; Papamichos-Chronakis *et al*. 2004; Roy *et al*. 2014). Tup1-Cyc8 remains at these promoters and aids in the recruitment of complexes such as SAGA and SWI/SNF that promote gene activation. Different sets of genes are controlled by distinct DNA-binding factors and are responsive to different kinases, illustrating that each pathway has developed its unique method of alternating between repressive and activating states.

A similar mechanism for corepressor to coactivator conversion may occur at genes regulated by α2-Tup1. At the *STE2* and *STE6* **a**-specific promoters, Tup1-Cyc8 remains bound upon removal of α2 and is important for Gcn5-dependent acetylation and rapid de-repression (Desimone and Laney 2010), suggesting Tup1 can act as a coactivator at in the absence of α2. Our observation of Tup1 occupancy at the *MFA2* promoter in both **a** and α cells is consistent with this hypothesis. In the α cell, α2-Tup1 interaction would repress expression of *MFA2*, but in the **a** cell, the absence of α2 would allow Tup1 to exist in its coactivator form, promoting expression of *MFA2*. Based upon previous studies, the position of the *tup1(S649F)* mutation could be expected to decrease or eliminate interaction with the α2 repressor protein (Komachi and Johnson 1997). The simultaneous increase in both *MFA2* expression and Tup1(S649F) occupancy upstream of *MFA2* in both **a** and α *tup1(S649F)* cells is therefore also consistent with this model (Fig S7). Likewise, the previous observation that a WD domain deletion mutant of Tup1 had a stronger de-repressive effect on *MFA2* than a *tup1* null (Komachi and Johnson 1997; Zhang *et al*. 2002) can be explained by a dual corepressor/coactivator function at *MFA2*. A *tup1* null constitutes a loss of both functions, whereas the disrupted α2 interaction caused by WD deletion or S649F mutation might affect only corepressor function. Since these mutants appear to retain coactivator capacity, they would lead to higher expression of target genes than a *tup1* null.

### Genomic redistribution of Tup1 occupancy in the *tup1(S649F)* mutant supports a change in the protein(s) that retain Tup1 at promoters

The types of genome-wide changes in Tup1 occupancy observed in the *tup1(S649F)* mutant relative to wild-type and the wide variety of genes with altered expression suggest that this mutation affects association of Tup1 not only with α2 but with additional DNA-binding proteins (Table S4, S5). Our ChIP-Seq data demonstrated reduction in Tup1 occupancy at some locations in the *tup1(S649F)* mutant, and a gain at even more locations (Fig 6B, C). At a number of sites, Tup1 had barely detectable association in wild-type but substantial occupancy in the mutant, and some peaks shifted relative to their original location (Fig S3A). Since Tup1 does not associate directly with DNA, these alterations imply a change in affinity of Tup1(S649F) for the proteins that tether it to DNA. The *tup1(S649F)* mutation could specifically affect the relevant WD domain binding interface necessary for these interactions or could affect association of more N-terminal repression domains with DNA-binding proteins due to a general loss of protein stability. The *tup1(S649F)* mutant could therefore represent on a genome-wide scale what has been observed previously at individual genes, that reduction or loss of interaction with the DNA-binding recruiter shifts Tup1 to a coactivator form but does not eliminate its association with chromatin. Our observation of substantially less Tup1 occupancy upstream of *MFA2* at the repressed gene (α cell) than at the active gene (**a** cell) also raises the possibility that interaction with DNA-binding proteins such as α2 could dampen the amount of Tup1 at the promoter. If so, this may be another factor contributing to locations of increased occupancy of Tup1(S649F) across the genome.

The most perplexing aspect of this model is the mechanism by which Tup1 is retained at promoters to facilitate gene activation in the absence of the interaction with the original DNA-binding recruiter protein. The answer may lie in the observation that Tup1 localizes within wide nucleosome depleted regions (NDRs) that have binding sites for many factors (Rizzo *et al*. 2011; Parnell *et al*. 2020), and promoters often show redundancy of recruitment of Tup1 by multiple DNA-binding proteins (Hanlon *et al*. 2011). In this way, Tup1-regulated gene promoters behave more like higher eukaryotic enhancers than most yeast promoters, and are capable of responding to input from multiple cellular conditions simultaneously (Hanlon *et al*. 2011; Rizzo *et al*. 2011). Loss of interaction with one DNA-binding protein may be critical for the switch to a coactivator state, but another could ensure retention of Tup1-Cyc8 at the promoter. Alternatively, once recruited to a promoter, Tup1 could remain associated by virtue of interaction with a more general protein, such as histones or the HMG-containing protein Nhp6, both shown to bind to Tup1 and/or Cyc8 under certain conditions (Laser *et al*. 2000; Fragiadakis *et al*. 2004). Given the large number of proteins with which Tup1 and Cyc8 are capable of forming interactions, including histone deacetylases and components of the Mediator complex (Kuchin and Carlson 1998; Gromoller and Lehming 2000; Papamichos-Chronakis *et al*. 2000; Watson *et al*. 2000; Han *et al*. 2001; Davie *et al*. 2003), many other scenarios are possible. This complexity highlights the importance of studying genes in their natural context and the limitations of reporter-based experiments in which transcription factor binding sites are isolated from other factors that normally influence their interaction with complexes such as Tup1-Cyc8.

### The role of Tup1 in regulation of mating type specific RNA expression extends beyond repression of a-specific genes

Our results question the simplicity of the long-standing model that Tup1-Cyc8 association with α2/Mcm1 at **a**-specific promoters causes their repression in α cells. Despite the dramatic differences in **a**-specific gene expression between **a** and α cells and between wild-type and *tup1(S649F)* mutant α cells, most promoters for these genes lacked substantial Tup1 occupancy or did not display the expected differential in Tup1 occupancy between **a** and a cells (Fig 8, S7). Most surprisingly, we observed strong association of Tup1 with the *MFA2* promoter within the **a** cell in which the gene is expressed, suggesting Tup1 could participate in activation as well as repression of *MFA2* within different cellular contexts. Another **a**-specific gene, *STE6*, requires Tup1 for full expression (Desimone and Laney 2010), raising the possibility that a coactivator function of Tup1 is required for activation of some of the **a**-specific genes.

While Tup1 has not previously been suggested to regulate α-specific genes, we observed Tup1 occupancy within the promoters of these genes at levels similar to or higher than those at the **a**-specific genes (Fig 8, S7). Similar to *MFA2*, strong Tup1 association was observed upstream of the α-specific gene *MF(ALPHA)1* within the α cell in which the gene is expressed, again suggesting a role for Tup1 in gene activation. Consistent with this idea, the group of α-specific genes, including *MF(ALPHA)1*, collectively display diminished expression in a *tup1* null (Chen *et al*. 2013). The promoters of *MF(ALPHA)2*, *SAG1* and *BSC1/PRM7* exhibit increased occupancy of Tup1(S649F) as well as increased expression in the α strain or in both mating types, further supporting a positive correlation between Tup1 occupancy and expression of the α-specific genes.

Our results therefore present a perplexing conundrum regarding the nature of Tup1 regulation of the mating type-specific genes, as upstream Tup1 occupancy did not necessarily predict its effects on transcription. In some cases, association of Tup1 appears to argue for an activator role of Tup1, but at other genes occupancy does not necessarily correlate with expression levels, particularly when considering repression of the **a**-specific genes. The strongest transcriptional effects of the *tup1(S649F)* mutant were observed at these **a**-specific genes, yet occupancy of Tup1 was not substantially altered. Perhaps some of the effects of Tup1 on expression could occur indirectly via its regulation of other transcription factors, a number of which display Tup1 occupancy within their upstream regions. Given the complex nature of promoters regulated by Tup1, additional as yet uncharacterized factors at these locations likely act in conjunction with or in opposition to Tup1, making it difficult to discern the direct relationship between Tup1 occupancy and expression levels. Nevertheless, the results presented here highlight that Tup1 likely plays a dual role in regulation of mating type in yeast, having the capacity to function as either a coactivator or a corepressor at different genes or in different mating types. The role of Tup1 in regulation of mating-type specific genes is therefore both more complex and more comprehensive than previously appreciated, and much remains to be learned about the nature of how Tup1-Cyc8 affects transcriptional processes.

## METHODS

### Strain and plasmid construction

Yeast strains are listed in Table S1 and are isogenic in the W303 background (*leu2-3,112 trp1-1 can1-100 ura3-1 ade2-1 his3-11,15*) (Thomas and Rothstein 1989). Standard strain construction methods were used (Rothstein 1991; Sherman 1991; Knop *et al*. 1999; Storici *et al*. 2001). Further details concerning plasmid and strain construction are available upon request.

For the genetic screens, *tup1* null strains were constructed by PCR amplification of either *HIS3MX* from pFA6:*HIS3MX6* (Longtine *et al*. 1998) or *URA3:KanMX* from pCORE with *Kluvermyces lactis URA3* and *KanMX4* (Storici *et al*. 2001), followed by integration at the *TUP1* locus to replace the ORF [*tup1(Δ+1 to +2142)*].

A *TUP1* YCp *TRP1* plasmid (M5765) was constructed by inserting a 3.5 kb PstI-HindIII fragment from plasmid pFW45 (Williams and Trumbly 1990) containing the *TUP1* ORF as well as 237 bp upstream and 1074 bp downstream endogenous sequence into YCplac22 *TRP1* (Gietz and Sugino 1988). To facilitate construction of *tup1* mutant alleles on plasmids, a pUC57 plasmid containing wild-type *TUP1* with three translationally silent restriction sites (SmaI, XmaI at amino acids 585-587, NruI at amino acids 636-637, and EcoRI at amino acids 675-677) was synthesized by Genewiz. PCR amplified fragments containing the relevant portions of mutant *tup1* plasmids isolated from genetic screens were cloned into the *TUP1* pUC57 vector and then used to replace wild-type *TUP1* in the *TUP1* YCp plasmid.

Endogenous *tup1* alleles were constructed by replacement of the *URA3(K.l.)::KanMX* cassette in DY18163 (*MAT***a** *tup1(Δ+1 to +2142)::URA3(K.l.)::KanMX*) with a PstI-HindIII fragment containing *TUP1* and flanking sequences from plasmids containing the relevant alleles. The mutant alleles inserted at the native *TUP1* locus were verified by PCR analysis and by sequencing. These *tup1* endogenous alleles were unmarked and were subsequently followed in crosses by qPCR, detecting the Tm difference between wild-type and mutant products from primers spanning the introduced silent EcoRV restriction site in mutant alleles (Wittwer *et al*. 2003).

The *TUP1-V5* and *tup1(S649F)-V5* alleles were constructed as described (Knop *et al*. 1999) by PCR amplifying and C-terminally integrating a V5 epitope tag with a *HIS3MX* marker from pZC03 (pFA6a-TEV-6xGly-V5-HIS3MX), provided by Zaily Connell and Tim Formosa (plasmid #44073; Addgene). Tagging of *TUP1* imparted a slight flocculation phenotype, and tagging of *tup1(S649F)* increased the severity of its flocculation phenotype.

### Genetic screens

Genetic screens for *tup1* mutants were conducted in *swi5 ash1* and *gcn5* mutant backgrounds using strains DY18884 (*MAT*α *HO-ADE2 HO-CAN1 swi5::KanMX ash1::LEU2 ade2::HphMX tup1(Δ+1 to +2142)::HIS3MX*) and DY19135 (MAT**a** HO-ADE2 HO-CAN1 gcn5::HIS3 ade2::HphMX tup1(Δ+1 to +2142)::URA3(K.l.)::KanMX), in which the *TUP1* ORF was replaced with a marker cassette. Strains were co-transformed with a mutagenized *TUP1* PCR product, whose primers introduced complementarity to the co-transformed linearized YCplac22 vector, allowing formation of complete plasmids containing *tup1* mutations via homologous recombination (Muhlrad *et al*. 1992), and mutants were selected on SD-Ade-Trp plates. For each screen, multiple *TUP1* PCR reactions were transformed and plated separately to allow for later determination of the independence of mutant alleles. Transformation of a wild-type *TUP1* plasmid (M5765) was always performed and plated separately for comparison purposes, as restoration of wild-type *TUP1* function from the plasmid allowed for better growth and a weak Ade+ phenotype relative to the starting *tup1* null strain. Seven candidate mutants were selected from each transformation, often of different sizes and ranging in color from pink to white, as *ADE2* expression affects colony color. All were then re-tested for growth on SD-Ade-Trp in comparison with a wild-type *TUP1* plasmid.

Candidate mutants from the *swi5 ash1* screen were initially verified by isolation of the mutant *tup1* plasmid, followed by re-transformation into the original *swi5 ash1 tup1 HO-ADE2* strain and re-testing of the growth phenotype on SD-Ade-Trp media. For those that maintained an Ade+ phenotype, plasmid DNA was isolated and subjected to Sanger sequencing using a series of primers that covered the *TUP1* gene. A large proportion of mutants that were re-tested displayed strong growth, demonstrating that the screen was very effective, with a low false positivity rate. Therefore, when performing the screen in the *gcn5* mutant, we quickly narrowed our interest to those mutant positions that appeared in multiple *tup1* plasmids or in both screens without re-testing each to confirm its phenotype. The extent to which the mutants in Table 1 were verified is indicated by their “Classification”. Many additional mutants were obtained from both screens but are not listed if they were observed in only one plasmid or were not independently re-tested for increased *HO-ADE2* expression. These include, but are not limited to: C348R (*gcn5* screen), I676M (*swi5 ash1* screen) and Y580H (*swi5 ash1* screen); these positions are noted due to their correspondence with mutations identified by Komachi et al (1997) that affected interaction of Tup1 with α2.

### Growth assays

For plate spot dilution assays, liquid cultures of the indicated strains were grown to saturation, serially diluted in 10-fold increments, spotted onto YPAD media for the number of days indicated, and photographed.

### Immunoblot analysis

Cells were grown at 30°C in YPA medium (1% yeast extract, 2% bactopeptone, 0.002% adenine) supplemented with 2% dextrose (Sherman 1991) to an OD_660_ of 1.0 to 1.5. For each sample, 1.0 OD of cells was collected, and protein lysate was prepared by pulverizing cells with glass beads in sodium dodecyl sulfate (SDS) buffer. Standard methods for sodium dodecyl sulfate-polyacrylamide gel electrophoresis (SDS-PAGE) were used. A nitrocellulose membrane (Bio-Rad) was used for western transfer with standard methods. Immunoblots were incubated with a 1:5,000 dilution of mouse monoclonal antibody to the V5 epitope (SV5-Pk1; Abcam) or a 1:10,000 dilution of mouse monoclonal antibody to Pgk1 (Invitrogen), followed by a 1:10,000 dilution of sheep anti-mouse antibody conjugated to horseradish peroxidase (HRP) (GE Biosciences). Antibody signals were detected with Super Signal West Dura Chemiluminescent Substrate (Thermo Scientific).

### RNA expression and chromatin immunoprecipitation (ChIP) analysis

Cells for RNA isolation or chromatin immunoprecipitation were grown at 30°C in YPA medium (1% yeast extract, 2% bactopeptone, 0.002% adenine) supplemented with 2% dextrose (Sherman 1991) to an OD_660_ of 0.6 to 0.8.

RNA was isolated and *HO* mRNA levels were measured by reverse transcription quantitative PCR (RT-qPCR), as described previously (Voth *et al*. 2007). *HO* RNA expression was normalized to that of *RPR1*, the RNA component of RNase P, transcribed by RNA polymerase III. *RPR1* transcription is not usually affected by genetic manipulations that alter RNA Pol II transcription.

ChIPs were performed as described (Bhoite *et al*. 2001; Voth *et al*. 2007), using mouse monoclonal antibody to the V5 epitope (SV5-Pk1; Abcam) and antibody coated magnetic beads (Pan Mouse IgG beads; Life Technologies). Cells were cross-linked in 1% formaldehyde overnight at 4°C and quenched with 125mM glycine. For ChIP qPCR experiments in Fig 6A, the concentration of ChIP DNA at the relevant target gene was normalized to its corresponding Input DNA and a No Tag control.

Quantitative PCR (qPCR) experiments for RNA expression and ChIP analysis were conducted on a Roche Lightcycler 480 or a ThermoFisher QuantStudio 3.

Concentrations were determined using wild-type complementary DNA (cDNA) or input ChIP DNA for in-run standard curves via the E-method (Tellmann 2006). The Student’s t-test was used to determine significance levels, as reported in the figure legends. Primers used for RT-qPCR and ChIP qPCR are listed in Table S2.

### RNA-Seq analysis

RNA was prepared as described (Ausubel 1987) and column purified using the QIAGEN RNeasy Mini Kit. Libraries were prepared for triplicate samples from each strain using the Illumina TruSeq Stranded Total RNA Ribo-zero Gold Library Prep reagent kit. Sequencing was performed on an Illumina NovaSeq 6000 as 50 bp paired-end runs (University of Utah High Throughput Genomics Facility). Fastq files were de-duplicated of optical duplicates using BBmap clumpify (version 38.34) and optical distance of 10000, trimmed of adapters using Cutadapt (version 2.8) with minimum overlap length (option -O) of 6 and minimum output length (option -m) of 20, and aligned to the genome (UCSC sacCer3) using the STAR aligner (version 2.7.3) with basic two-pass mode. Read counts over transcripts were collected using subRead featureCounts (version 1.6.3) using custom transcript annotation that includes UTRs and excludes dubious annotation (Parnell 2021b). Differential analysis was performed with DESeq2 with the aid of the hciR package (Parnell 2021a). Differential genes that were selected by Custom heat maps were made with pHeatmap and volcano plots with ggPlot in custom R scripts.

Only protein-coding genes were considered for quantitation of genes that demonstrated changes in gene expression (Fig 5A), and any with a base mean of less than 30 were eliminated. Genes counted in Fig 5A had a log_2_ fold expression change of greater than +1.0 or less than −1.0 (2-fold change). For the Tup1 occupancy ChIP-Seq vs. RNA expression RNA-Seq correlations in Fig 7B, 7D, and S4-6, any changes greater than +0.585 or less than −0.585 (1.5-fold change) were counted.

### ChIP-Seq analysis

Chromatin isolated from individual, independently collected Tup1-V5 or Tup1(S649F)-V5 cell pellets was used for multiple ChIPs, performed as described above, which were then pooled for each replicate. Libraries were prepared for triplicate ChIP samples and a single input sample for each strain using the New England Biolabs NEBNext ChIP-Seq Library Prep Reagent Set with dual index primers. Sequencing was performed with an Illumina NovaSeq 6000, 50 bp paired-end run in two lanes (University of Utah High Throughput Genomics Facility). Fastq files were aligned to the genome (UCSC sacCer3) using Novocraft Novoalign version 4.2.2, giving primer adapters for trimming, Novaseq tuned parameters, and allowing for 1 random, multi-hit alignment. Alignments from individual sequencing runs were merged for each sample. An average of 13 million fragments were mapped.

Samples were then processed with MultiRepMacsChIPSeq pipeline version 14.1 (Parnell 2020). Alignments over mitochondrial, 2-micron, rDNA, and telomeric regions were discarded from analysis. Excessive duplicate alignments (range 7-16%) were randomly subsampled to a uniform 6% for each sample after removing optical duplicates (minimum pixel distance of 10000). Replicates were depth-normalized, averaged together, and peak calls generated with a minimum length of 250 bp, gap size of 50 bp, and minimum q-value statistic of 3. Alignment counts from each replicate was collected for each peak and used for correlation metrics and identifying differentially occupied peaks by running DESeq2, version 1.28 (Love *et al*. 2014). Peaks were annotated by intersection using bedtools (Quinlan 2020) with interval files of either genes or intergenic regions. Heat maps were generated by first collecting data with BioToolBox get_relative_data from the peak midpoint 25 windows of 25 bp, and plotting the data using pHeatmap (Kolde 2020) in custom R scripts.

Peaks that were annotated as “Intergenic”, as described above, with a flanking feature of a tRNA, other Pol III gene (*SCR1*, *RPR1*) or snRNA, were individually examined in the IGV genome browser (Broad Institute) to determine whether the peak overlapped the tRNA/snRNA or whether the peak was located in the intergenic region between a tRNA/snRNA and another gene. Peaks overlapping tRNA/snRNA genes were classified as “Poll III, snRNA” and the peaks between tRNA/snRNA and/or Pol II regulated genes were grouped as “Intergenic” peaks, and these designations were used in Fig S3A and Table S7. Some of the “UTR/ORF” peaks in Fig S3A had Tup1 occupancy that spanned intergenic and UTR sequences upstream of a possible target gene. For Table 2 analysis, these peaks were classified as “Upstream”, based upon examination in the IGV genome browser (Broad Institute).

### Data availability

Strains and plasmids are available upon request. Table S1 lists the strains used in this study. Primers used for RT-qPCR and ChIP qPCR analysis are listed in Table S2. RNA-Seq differential expression data is shown in Table S3; genes with >2-fold change are listed in Table S4, along with the cluster in which they were placed (Fig 5B). All ChIP-Seq peaks with location and cluster information (Fig 7A) are listed in Table S7; flanking genes associated with each are also indicated. RNA-Seq and ChIP-Seq data have been deposited in NCBI’s Gene Expression Omnibus with the accession number GSE173722.

## ACKNOWLEDGEMENTS

We thank Tim Formosa and members of the Stillman lab for advice throughout the course of these experiments and Tim Formosa for comments on the manuscript. We thank high school summer students Micah Spjute and Ellie Siddoway for help with genetic screens for *tup1* mutants. Plasmid pFW45 containing the *TUP1* gene was provided by Robert Trumbly, and plasmid pZC03 (pFA6a-TEV-6xGly-V5-HIS3MX) was provided by Zaily Connell and Tim Formosa. This work was supported by National Institutes of Health grant GM-121079 awarded to D.J.S.

---

**Figure S1. Comparison of Tup1 vs. Tup1(S649F) protein levels in a vs. α strains.** Relative protein levels for Tup1-V5 and Tup1(S649F)-V5 strains were determined by western immunoblot analysis. Total yeast lysate from equal numbers of cells, as judged by optical density, were loaded for each lane. Anti-V5 antibody (Abcam, SV5-Pk1) was used for the top portion of the blot, and anti-Pgk1 (Invitrogen) was used for the bottom portion of the blot as a loading control.

**Figure S2. Pearson correlation plot for ChIP-Seq of Tup1-V5 and Tup1(S649F)-V5 in a and α strains.** Shown is a genome-wide Pearson correlation plot, using ChIP-Seq fragment counts in bins of 1 kb. Each strain is represented by triplicate ChIP samples and a single Input sample. Hierarchical clustering displays the relationships between samples; color coding for clustering of ChIP and Input samples is shown at the right.

**Figure S3. Categorization and occupancy features of genomic Tup1 peaks.** (A) Genome snapshots from the genome browser IGV (Broad Institute) show ChIP-Seq occupancy results for a portion of chromosome XI with representative peaks that seem to appear or shift position in the *tup1(S649F)* mutant strains. Each track represents the average of triplicate samples. Colors are as follows: Tup1-V5 WT **a** (red), Tup1-V5 WT α (green), Tup1(S649F)-V5 **a** (blue) and Tup1(S649F)-V5 α (purple). The bottom track displays gene annotations. Both panels show the same genomic location; the top panel is scaled differently than the lower panel to illustrate Tup1 enrichment in strains with vastly different occupancy levels. (B) The table lists the number and percentage of Tup1 peaks in different genomic locations: intergenic, UTR/ORF, or Pol III / snRNA depending upon the position of the majority of peak occupancy. (C) Genome snapshots from the genome browser IGV (Broad Institute) show ChIP-Seq occupancy results for a portion of chromosome II that contains a tRNA peak and an intergenic double peak. Each track represents the average of triplicate samples. Colors are as follows: Tup1-V5 WT **a** (red), Tup1-V5 WT α (green), Tup1(S649F)-V5 **a** (blue) and Tup1(S649F)-V5 α (purple). The bottom track displays gene annotations. Both panels show the same genomic location; the top panel is scaled differently than the lower panel to illustrate the difference in Tup1 enrichment between the intergenic and tRNA peaks. (D) Violin plots display the log_2_ fold change in Tup1(S649F) to Tup1 wild-type occupancy in **a** and α strains at groups of peaks, including those located within intergenic regions (Left), over UTR/ORFs (Middle) and over Pol III genes and snRNAs (Right). Each point represents a single peak from the Table in (A).

**Figure S4. Scatter plots to illustrate Tup1 occupancy vs. RNA expression in *tup1(S649F)* vs. wild-type a strains.** The log_2_-fold change in ChIP occupancy between Tup1(S649F) and wild-type Tup1 **a** strains was plotted relative to the log_2_-fold change in RNA-Seq expression levels between *tup1(S649F)* and wild-type. Each dot represents a unique gene / peak combination. The sides of the red squares at the graph origins demarcate 1.5-fold changes in occupancy or expression (log_2_ = 0.585 or −0.585); points located within the box are those scored as no change, “NC”. Tup1 peaks located between divergent genes were used twice in the analysis, once for each gene. Some genes were counted more than once, due to the presence of multiple Tup1 peaks within their upstream intergenic region (53 genes counted as 118 out of 1039 total genes). In most of these cases, the peaks fall in different cluster categories.

**Figure S5. Scatter plots to illustrate Tup1 occupancy vs. RNA expression in *tup1(S649F)* vs. wild-type α strains.** The log_2_-fold change in ChIP occupancy between Tup1(S649F) and wild-type Tup1 α strains was plotted relative to the log_2_-fold change in RNA-Seq expression levels between *tup1(S649F)* and wild-type. Each dot represents a unique gene / peak combination. The sides of the red squares at the graph origins demarcate 1.5-fold changes in occupancy or expression (log_2_ = 0.585 or −0.585); points located within the box are those scored as no change, “NC”. Tup1 peaks located between divergent genes were used twice in the analysis, once for each gene. Some genes were counted more than once, due to the presence of multiple Tup1 peaks within their upstream intergenic region (53 genes counted as 118 out of 1039 total genes). In most of these cases, the peaks fall in different cluster categories.

**Figure S6. Heat maps to illustrate Tup1 occupancy vs. RNA expression in *tup1(S649F)* vs. wild-type a strains.** Shown is the Tup1 occupancy vs. RNA expression data for clusters of Tup1 peaks in **a** strains, analyzed in the same manner as the α strains in Fig 7B. Each square within the 3 x 3 heat maps represents a unique combination of Tup1 occupancy and RNA expression of the downstream gene, where “Up” and “Down” indicate increased or decreased occupancy/expression in *tup1(S649F)* α vs. wild-type α, respectively, and “NC” (No Change) indicates less than 1.5-fold alteration in levels. The color scale at the right indicates the percentage of genes in each category. Genes were separated by cluster from Fig 7A and were only included if they were located downstream of a Tup1 peak and had a valid expression level from RNA-Seq (993 total). Peaks were counted twice if present between divergent genes. Some genes were counted more than once, due to two or three Tup1 peaks within the intergenic region upstream of the gene (53 total).

**Figure S7. Genes that are differentially expressed in a vs. α strains.** IGV genome browser (Broad Institute) snapshots of ChIP-Seq (Occupancy) and RNA-Seq (Expression) show the sequenced fragment pileups for Tup1 peaks located next to the thirteen mating-type specific genes. Tracks represent an average of three biological samples for each strain. The four strains for each experiment were autoscaled as a group for occupancy and expression separately to account for differences in fragment depth between locations. Colors for both ChIP-Seq and RNA-Seq are as follows: Tup1-V5 WT **a** (red), Tup1-V5 WT α (green), Tup1(S649F)-V5 **a** (blue) and Tup1(S649F)-V5 α (purple). The bottom track displays gene annotations.

